# Redox-Dependent Condensation Of the Mycobacterial Nucleoid By WhiB4

**DOI:** 10.1101/133181

**Authors:** Manbeena Chawla, Saurabh Mishra, Pankti Parikh, Mansi Mehta, Prashant Shukla, Manika Vij, Parul Singh, Kishore Jakkala, H N Verma, Parthasarathi AjitKumar, Munia Ganguli, Aswin Sai Narain Seshasayee, Amit Singh

**Author notes:** These authors contributed equally to this work (Manbeena Chawla, Saurabh Mishra). Correspondence: Amit Singh, Amit Singh, Department of Microbiology and Cell Biology (MCBL), Centre for Infectious Disease and Research (CIDR), Indian Institute of Science (IISc), Bangalore, India-560012.

## Abstract

Oxidative stress response in bacteria is generally mediated through coordination between the regulators of oxidant-remediation systems (*e.g.* OxyR, SoxR) and nucleoid condensation (*e.g.* Dps, Fis). However, these genetic factors are either absent or rendered nonfunctional in the human pathogen *Mycobacterium tuberculosis* (*Mtb*). Therefore, how *Mtb* organizes genome architecture and regulates gene expression to counterbalance oxidative imbalance during infection is not known. Here, we report that an intracellular redox-sensor, WhiB4, dynamically links genome condensation and oxidative stress response in *Mtb*. Disruption of WhiB4 affects the expression of genes involved in maintaining redox homeostasis, central carbon metabolism (CCM), respiration, cell wall biogenesis, DNA repair and protein quality control under oxidative stress. Notably, disulfide-linked oligomerization of WhiB4 in response to oxidative stress activates the protein’s ability to condense DNA *in vitro* and *in vivo*. Further, overexpression of WhiB4 led to hypercondensation of nucleoids, redox imbalance and increased susceptibility to oxidative stress, whereas WhiB4 disruption reversed this effect. In accordance with the findings *in vitro*, ChIP-Seq data demonstrated non-specific binding of WhiB4 to GC-rich regions of the *Mtb* genome. Lastly, data indicate that WhiB4 deletion affected the expression of only a fraction of genes preferentially bound by the protein, suggesting its indirect effect on gene expression. We propose that WhiB4 is a novel redox-dependent nucleoid condensing protein that structurally couples *Mtb’s* response to oxidative stress with genome organization and transcription.

**Significance Statement:** *Mycobacterium tuberculosis (Mtb)* needs to adapt in response to oxidative stress encountered inside human phagocytes. In other bacteria, condensation state of nucleoids modulates gene expression to coordinate oxidative stress response. However, this relation remains elusive in *Mtb*. We performed molecular dissection of a mechanism controlled by an intracellular redox sensor, WhiB4, in organizing both chromosomal structure and selective expression of adaptive traits to counter oxidative stress in *Mtb*. Using high-resolution sequencing, transcriptomics, imaging, and redox biosensor, we describe how WhiB4 modulates nucleoid condensation, global gene expression, and redox-homeostasis. WhiB4 over-expression hypercondensed nucleoids and perturbed redox homeostasis whereas WhiB4 disruption had an opposite effect. Our study discovered an empirical role for WhiB4 in integrating redox signals with nucleoid condensation in *Mtb*.

## Introduction

*Mycobacterium tuberculosis* (*Mtb*) is the causative agent of tuberculosis (TB), which is one of the major global human health problems. Upon aerosol infection, *Mtb* is phagocytosed by alveolar macrophages, where it is exposed to different hostile conditions like reactive oxygen and nitrogen species (ROS and RNS), nutrient starvation, acidic pH and antimicrobial peptides (1, 2). Among the host responses, the contribution of redox stress in controlling the proliferation of *Mtb* has been studied in detail. For example, mice deficient in ROS generating enzyme, NADPH oxidase (NOX-2) showed greater susceptibility to *Mtb* infection (3). Studies demonstrating the increased susceptibility of children with defective NOX-2 to infections caused by *Mtb* and BCG further underscore the clinical relevance of the oxidative stress response (4). However, *Mtb* has several resistance mechanisms to overcome the redox stress it encounters inside macrophages. These include AhpC/D (alkyl hydroperoxidase), KatG (catalase), SodA and SodC (superoxide dismutases), TrxA, TrxB1 and TrxC (thioredoxins), glucose-6-phosphate dehydrogenase, MsrA/B (methionine-sulfoxide reductase), and the membrane-associated oxidoreductase complex (MRC) (5, 6). Furthermore, cytoplasmic antioxidants like mycothiol (MSH) and ergothioneine (ERG) are known to dissipate redox stress during infection (7, 8).

Apart from these response mechanisms, the bacterial nucleoid also undergoes dynamic changes in its architecture in response to stresses such as ROS, RNS, and changes in osmolarity and pH (9). Nucleoid-associated proteins (NAPs) including Dps (<u>D</u>NA-binding <u>p</u>rotein from <u>s</u>tarved cells) and Fis (<u>F</u>actor for <u>i</u>nversion <u>s</u>timulation) in *E. coli*, MrgA (<u>M</u>etallo <u>r</u>egulated <u>g</u>enes <u>A</u>) in *Staphylococcus aureus*, DR0199 (the homolog of EbfC) in *Deinococcus radiodurans*, HU (the homolog of Hup protein) in *Helicobacter pylori*, and Lsr2 in *Mtb* protect the cells from Fenton-mediated oxidative damage by physically shielding and compacting genomic DNA (10-14). Whereas regulated condensation of the nucleoid protects the cells against oxidative stress, long-lasting condensation has been shown to perturb normal cellular processes such as replication and transcription and stimulate oxidative stress-mediated death (15). Despite the recognized importance of bacterial NAPs in modulating the oxidative stress response, the effector proteins that monitor the changes in the cytoplasmic redox potential and sculpt the genome architecture in response to oxidative stress remain unknown. In *Mtb,* several studies have demonstrated the role of redox-responsive WhiB proteins (WhiB1 to WhiB7) in regulating gene expression and controlling a plethora of functions including antibiotic resistance, oxidative/nitrosative/acidic stress response, immune-response, cell division, and secretion (16-26). However, several functional aspects of WhiB proteins in *Mtb* have not been investigated. Previously, we have shown that one of the WhiB family member, WhiB4, is required to regulate survival in response to oxidative stress *in vitro* and to regulate virulence *in vivo* (22). A fundamentally important question remains yet unanswered; what is the mechanism by which WhiB4 coordinates oxidative stress response in *Mtb*? More importantly, while electrophoretic mobility shift assays (EMSAs) revealed DNA binding activity of some of the *Mtb* WhiB proteins [20,21,24,26], an *in vivo* evidence for redox-dependent DNA binding at a genomic scale using chromatin immunoprecipitation sequencing (ChIP-seq) is still lacking.

In this study, we employed multiple analytical techniques including microarray, atomic force microscopy (AFM), transmission electron microscopy (TEM), confocal microscopy, and ChIP-seq to examine the role of WhiB4 in modulating nucleoid condensation and oxidative stress response. Furthermore, using a non-invasive biosensor of the mycothiol redox potential (*E_MSH_*, Mrx1-roGFP2), we provide evidence for a functional linkage between redox physiology and genome compaction in *Mtb*. Our results show that in *Mtb* WhiB4 dynamically manipulates both DNA architecture and gene expression to control oxidative stress response during infection.

## Results

### *Mtb* WhiB4 regulates gene expression in response to oxidative stress

Previously, we had shown that WhiB4 functions as an autorepressor and marginally influences the expression of 23 genes in *Mtb* under unstressed conditions (22). Furthermore, a *whiB4* mutant survived better under oxidative stress *in vitro*, in immune-activated macrophages, and was hypervirulent in animals (22). In agreement with these findings, exposure to increasing concentrations of an oxidant, cumene hydroperoxide (CHP) induced a dose dependent decrease in WhiB4 expression as compared to that of 16s rRNA (Fig. 1A). These findings indicate that the expression of WhiB4 and its regulatory function thereof could be dependent on oxidative stress.

**Figure 1:**
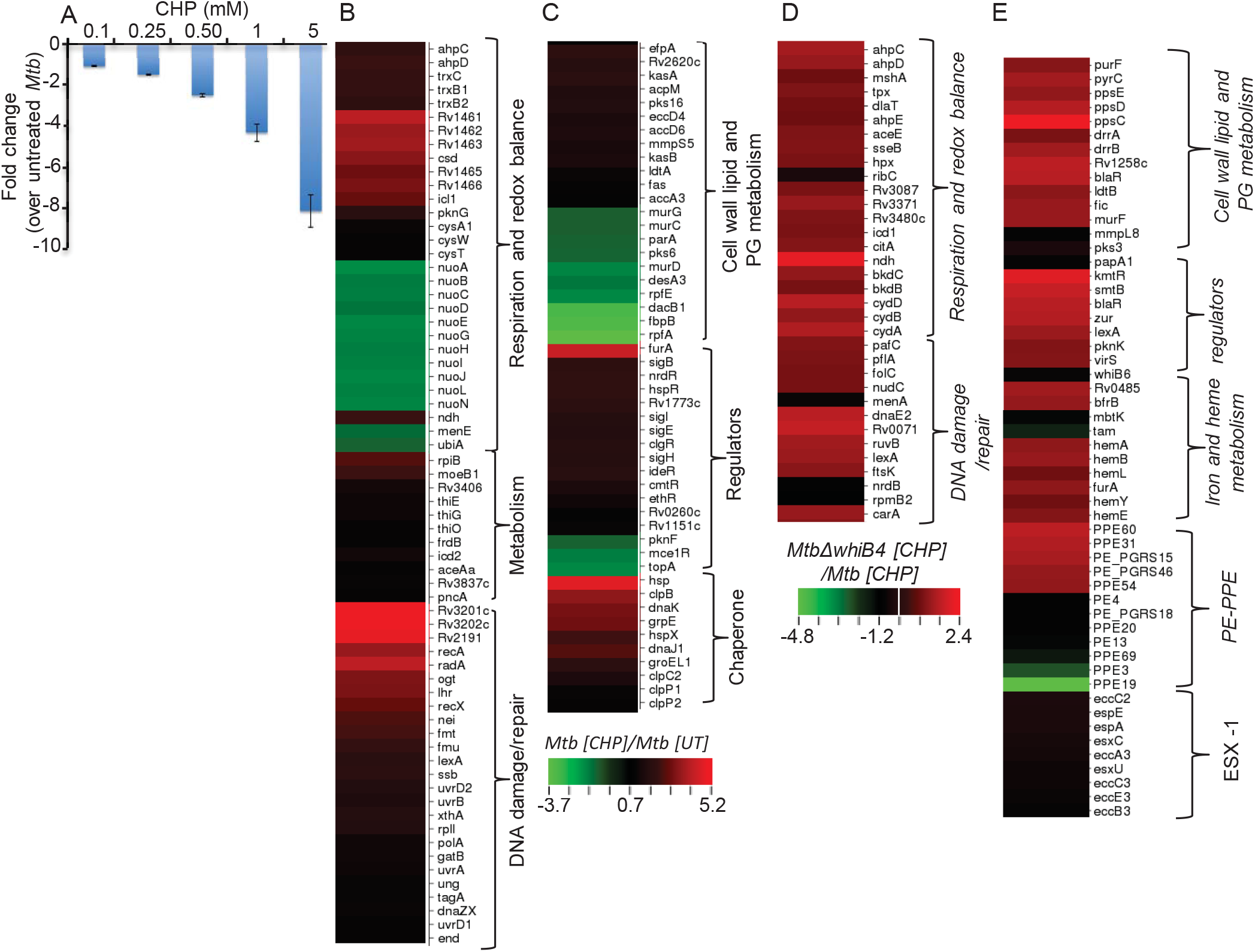
WhiB4 mediated regulation of gene expression in response to oxidative stress in *Mtb*. **(A)** RNA was isolated from wt *Mtb* upon treatment with the indicated concentrations of CHP for 2 h. qRT-PCR for *whiB4* was done using *whiB4*-specific oligonucleotides and fold change was normalized to 16s rRNA expression. **(B, C, D, and E)** Heat maps depicting the expression of genes (fold-change; *P* ≤0.05) coordinating respiration, CCM, DNA damage, PE-PPE and redox balance in response to 250 μM for 2 h of CHP-treatment of *Mtb* and *Mtb∆whiB4* from two biological samples.

Since the WhiB4 mutant reported earlier retained expression from initial 96 bases (22), we first generated an unmarked strain of *Mtb* (*MtbΔwhiB4*) lacking the entire open reading frame (ORF) encoding WhiB4 (Fig. S1). Next, we performed global transcriptome profiling of wt *Mtb* and *MtbΔwhiB4* upon exposure to 0.25 mM CHP. We minimized any influence of oxidative stress-induced cell death on gene expression by performing microarrays at an early time point (2 h) post-CHP treatment. In contrast to unstressed conditions, oxidative stress influenced the expression of a large number of genes in wt *Mtb* and *MtbΔwhiB4* in response to CHP [(log values) 1.5-fold up- and down-regulation, p≤0.05] (S1 Table).

First, we analyzed the effect of CHP on wt *Mtb*. Expectedly, genes directly implicated in mitigating oxidative stress were induced by CHP. This includes genes encoding thioredoxins, catalase, alkyl hydroperoxidase, and Fe-S biogenesis/repair operon (Fig. 1B and S1 Table). The eukaryotic-type protein kinase G (*pknG*) and the glyoxylate cycle enzyme isocitrate lyase (*icl*) have been recently shown to maintain redox homeostasis (27-29). Both the genes were significantly induced upon CHP treatment (Fig. 1B). Since oxidative stress damages DNA, protein, and lipids, a significant fraction of genes mediating DNA repair, cell wall lipids biogenesis, and protein quality control systems were induced by CHP (Fig. 1B, 1C). Our data shows that genes encoding energetically efficient respiratory complexes such as NADH dehydrogenase I (*nuo* operon), menaquinione (*menE*), and ubiquinone (*ubiA*) were down-regulated, whereas the energetically less favored NADH dehydrogenase type II (*ndh*), and fumarate reductase (*frdB*) were induced in response to CHP (Fig. 1B). These changes along with the induction of genes associated with the pentose phosphate pathway, glyoxylate shunt, and nicotinamide metabolism involved in maintaining NAD(P)H/NAD^+^(P) poise implicate a strategic shift from energy generation to redox balance in response to CHP stress. Importantly, oxidative stress triggers the expression of several transcription factors/sigma factors controlling iron homeostasis (*furA* and *ideR*), protein turn-over (*hspR*), redox homeostasis and cell envelope architecture (*clgR, ethR, nrdR, sigE, sigB,* and *sigH*) (Fig. 1C). In other bacterial species, dynamic changes in the nucleoid condensation state directed by several NAPs, and assisted by topology regulating enzymes such as DNA gyrases and topoisomerases, protects against oxidative stress (12, 30, 31). Surprisingly, microarray analysis showed that *topA* was down-regulated and the expression of NAPs (*espR, lsr2,* and *hupB*) was unaffected upon oxidative stress in *Mtb* (S1 Table).

Thereafter, we examined the role of WhiB4 in regulating oxidative stress-induced changes in gene expression. Approximately 280 genes were induced and 200 were repressed in *MtbΔwhiB4* as compared to that in *wt Mtb* upon CHP-treatment (>1.5-fold; p<0.05) (S1 Table). Our results revealed that genes playing overlapping roles in redox-metabolism and central carbon metabolism (CCM) are significantly induced in *MtbΔwhiB4*in response to CHP as compared to that in wt *Mtb* (Fig. 1D). This subset includes*, dlaT, bkdC*, *aceE*, *ahpE*, and *ahpCD*, which serve as functional subunits of the CCM enzymes - pyruvate dehydrogenase (PDH), α-ketoglutarate (α-KG) dehydrogenase, and branched-chain keto-acid dehydrogenase (BKADH). Most of these enzymatic activities are well known to confer protection against oxidative and nitrosative stress in *Mtb* (32-37). Moreover, similar to wt *Mtb*, the primary NADH dehydrogenase complex (*nuo operon*) was down-regulated in *MtbΔwhiB4* in response to CHP treatment (S1 Table). However, compensatory increase in the alternate respiratory complexes such as *ndh, frdA,* and cytochrome bd oxidase (*cydAB*) was notably higher in *MtbΔwhiB4* than in wt *Mtb*, indicating that *MtbΔwhiB4* is better fit to replenish reducing equivalents during CHP-induced cellular stress (Fig. 1D and S1 Table). In bacteria, including *Mtb*, cytochrome bd oxidase also displays catalase and/or quinol oxidase activity (38, 39), which confers protection against oxidative stress. Expression data is consistent with the greater potential of *MtbΔwhiB4* to tolerate oxidative stress, which is further supported by a substantially lesser number of DNA repair genes induced by CHP in *MtbΔwhiB4* as compared to *wt Mtb (*Fig. 1D). Several members of the PE_PGRS gene family and toxin-antitoxin modules, involved in maintaining cell wall architecture, protection from oxidative stresses, and drug-tolerance were found to be expressed at higher levels in *MtbΔwhiB4* (Fig. 1E and S1 Table) (40). Interestingly, while the deletion of WhiB4 increased the expression of transcription factors involved in metal sensing (*kmtR*, *smtB,* and *zur*), antibiotic tolerance (*blaR*), and virulence (*pknK-virS*), the expression of *whiB6* and *esx-1* secretion system was found reduced (Fig. 1E). Lastly, microarray data was validated by measuring the expression of a selected set of CHP- and *whiB4*-dependent genes by qRT-PCR (Table S2). Altogether, WhiB4 affects the expression of genes involved in oxidative stress response, alternate respiration, CCM, and PE_PGRS family in *Mtb*.

### *Mtb* WhiB4 possesses NAP-like DNA binding properties

We have previously shown that four cysteine residues in WhiB4 coordinate a 4Fe-4S cluster (holo-WhiB4), which can be rapidly degraded by atmospheric oxygen to generate clusterless apo-WhiB4 [26]. The exposed cysteine residues of apo- WhiB4 further oxidizes to generate disulfide-linked dimers and trimers of WhiB4 (22). Our earlier study on WhiB4 DNA binding at a specific locus (*ahpCD*) showed that holo-WhiB4 or reduced apo-WhiB4 lacks DNA binding capacity, whereas oxidized apo-WhiB4 binds at a specific locus (*ahpCD*) in a sequence-independent manner (22). These characteristics, along with the low molecular weight (13.1 kDa) and a highly basic pI (10.28) of WhiB4, are reminiscent of various nucleoid-associated proteins (NAP) (*e.g.,* HNS, HU, IHF and Lrp) (9). Since CHP stress is likely to generate DNA binding proficient form of WhiB4 (i.e. oxidized apo-WhiB4) *in vivo*, we hypothesized that under oxidative conditions WhiB4 influences gene expression by interacting non-specifically with the nucleoid. Using *in vivo* thiol-trapping experiment (*see Materials and Methods*), we confirmed that pretreatment with 0.1 and 0.5 mM of CHP significantly increased the proportion of disulfide-linked dimer (10,000-14,000 molecules) and trimer (4000-6000 molecules) of oxidized apo-WhiB4 per *Mtb* cell *(see SI Note I* and Fig. S2). In the next stage, we systematically investigated whether oxidized apo-WhiB4 possesses a NAP-like DNA binding properties *in vitro* and inside *Mtb*.

Similar to other bacterial NAPs (9, 11), the oxidized apo-WhiB4 formed a high molecular weight complex with a range of DNA substrates (*e.g.,* supercoiled DNA, linearized DNA, and 1kb λ DNA ladder) and inhibited transcription from a standard T7-promoter of pGEM plasmid *in vitro* (Fig. S3A-E). The inclusion of the thiol-reductant, dithiothreitol (DTT), or replacement of any cysteine residues with alanine in WhiB4, reversed the DNA binding and transcriptional inhibitory activities of oxidized apo-WhiB4 (Fig. S3C-D). Further, we examined apo-WhiB4 DNA binding properties using Atomic Force Microscopy (AFM). A super-coiled plasmid DNA (pEGFP-C1) was pre-exposed to increasing concentrations of oxidized apo-WhiB4 and subjected to AFM. The oxidized apo-WhiB4 and plasmid DNA were taken as the experimental controls. AFM images in the absence of WhiB4 showed uniform structure for the negatively supercoiled plasmid DNA (Fig. 2A). Interestingly, at lower amounts of oxidized apo-WhiB4, a gradual relaxation of supercoiled plasmid DNA with initial opening at the ends, followed by a mixed population of partially or fully opened DNA circles was detected (Fig. 2B-C). Further increase in oxidized apo-WhiB4 resulted in the formation of multiple loops, bends and extensive condensation within a single DNA molecule (Fig. 2D-3F). A further increase in the protein concentrations may lead to nucleoprotein filament formation. However, at higher concentrations of oxidized apo-WhiB4 forms large aggregates on the mica, thus precluding any further AFM imaging. An assessment of various geometric properties of the imaged protein:DNA complexes confirmed an initial increase in diameter and length followed by a decrease in both parameters at higher protein concentrations (Fig. 2G). In contrast, the Cys3-WhiB4 (replacement of third cysteine to alanine) mutant protein was unable to exhibit a similar effect on supercoiled pEGFP-C1 (Fig. S4), emphasizing the redox-dependent DNA condensing activity of WhiB4. This dual compactional ability of WhiB4 indicates a stoichiometry-dependent regulatory role for WhiB4, as shown for other chromosomal architectural proteins HU and RdgC in *E.coli* (41-43).

**Figure 2:**
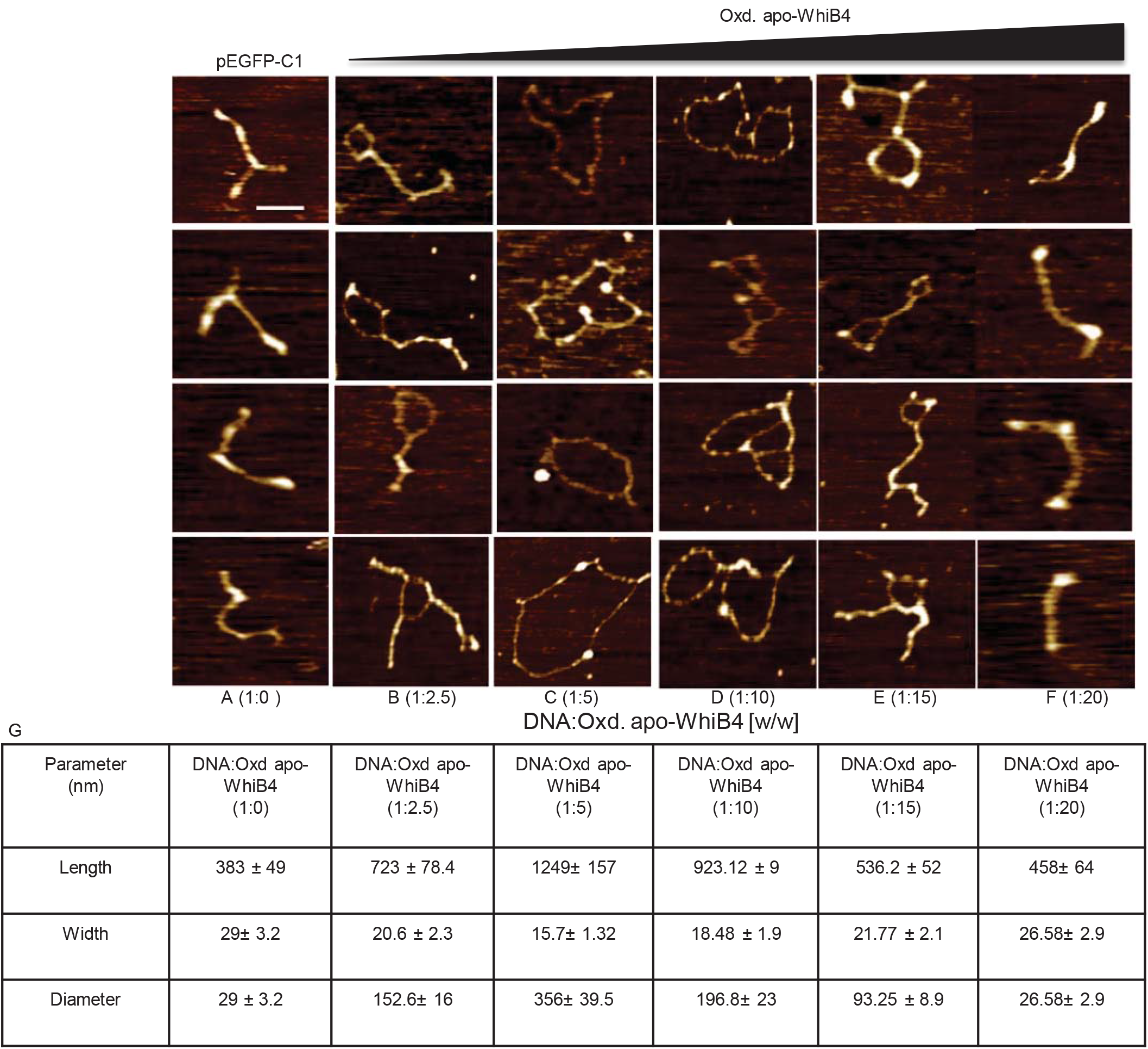
AFM analysis of WhiB4-mediated DNA condensation. **(A)** Supercoiled plasmid DNA, **(B-F)** Oxidized apo-WhiB4 incubated with supercoiled plasmid in increasing concentration (DNA: protein w/w =1:2.5, 1:5,1:10,1:15, and 1:20). **(B and C)** Low protein to DNA ratio gradually opens up the DNA until a fully relaxed circular DNA molecule is formed as observed in the case of a moderate protein to DNA ratio. **(D and E)** Further increase in protein to DNA ratio reverses the phenomenon when lateral compaction occurs. **(F)** High protein to DNA ratio further causes condensation of the DNA molecules thereby leading to rigid filamentous structures. The scale of images is 0.8 μm x 0.8 μm/ Scale bar is 100nm. **(G)** Geometric properties of the imaged protein: DNA complexes. Geometrical parameters of supercoiled DNA in the absence and presence of oxidized apo-WhiB4. (A- DNA only, B- low protein to DNA ratio, C-E- moderate protein to DNA ratio, F- high protein to DNA ratio; n= 70 independent DNA molecules measured in each case).

### WhiB4 condenses the mycobacterial nucleoid

In order to further understand the link between genome condensation, oxidative stress, and WhiB4, we examined the nucleoid morphology of wt *Mtb*, *MtbΔwhiB4*, *whiB4-Comp,* and *whiB4-OE* strains. The *whiB4-Comp* strain was generated by expressing *whiB4* under the native promoter in *MtbΔwhiB4*, whereas *whiB4-OE* strain allows overexpression of the FLAG-tagged *whiB4* from an anhydrotetracycline (Atc)-inducible promoter system in *MtbΔwhiB4*,. We have earlier shown that WhiB4 predominantly exists in an oxidized apo-form upon Atc-induced overexpression in a non-pathogenic fast growing *Mycobacterium smegmatis* (*Msm*) during aerobic growth (26). Since oxidized apo-WhiB4 acts as an autorepressor (26), continued expression of WhiB4 by Atc circumvents this regulatory system and likely amplifies the negative effect of WhiB4 on the expression of antioxidant systems. As a result oxidative stress increases inside *Mtb* leading to oxidation of apo-WhiB4 thiols and generation of oxidized apo-WhiB4 oligomers. Consistent with this, we confirmed that Atc-induced overexpression of WhiB4 resulted in a higher proportion of disulfide-linked dimeric and trimeric forms of oxidized apo-WhiB4 in *whiB4-OE* (*SI Note II* and Fig. S5). Quantification of intracellular abundance of WhiB4 revealed that the induction using 100 ng/ml of Atc generated 14,000 dimers and 4000 trimers of oxidized apo-WhiB4 per cell (*SI Note II* and Fig. S5), which is comparable to the oxidized apo-WhiB4 forms generated in a *Mtb* cell upon exposure to 0.1-0.5 mM CHP.

We stained the nucleoids of *wt Mtb*, *MtbΔwhiB4*, *whiB4-Comp* and *whiB4-OE* strains with 4’,6-diamidino-2-phenylindole (DAPI) and visualized the cells by confocal microscopy (see SI Materials and Methods). DAPI is extensively used to illuminate shape and size of nucleoids in a range of bacteria including *Mtb* (44, 45). DAPI stained cells from exponentially grown cultures of *Mtb* strains frequently showed the presence of expanded nucleoids with a fewer bilobed nucleoid. Infrequently, single cells containing more than 3 distinct DAPI stained regions were also observed (Fig. 3). Using these images we measured relative nucleoid size (RNS) by determining the ratio between the length of the nucleoid(s) and the length of the cell by relying on the end points of their larger axes. Since the images are two-dimensional and cells are small, the values of RNS are merely approximations indicating the trend of compactions or expansion of the nucleoid under conditions studied.

**Figure 3:**
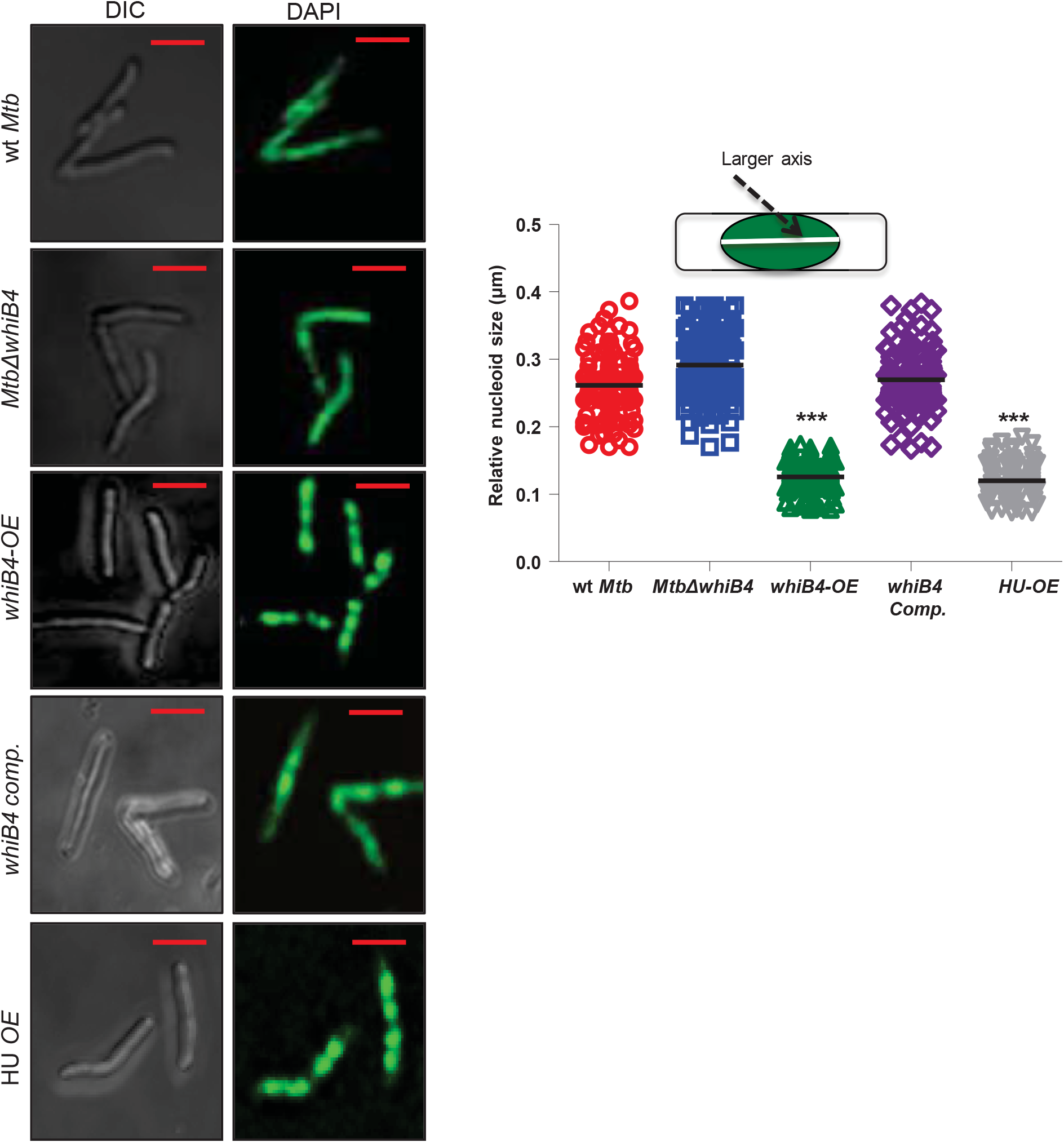
WhiB4-mediated nucleoid compaction in *Mtb*. **(A)** Nucleoid of wt *Mtb*, *MtbΔWhiB4*, *whiB4-OE, whiB4 comp.* and *HU OE* strains were stained with DAPI (pseudocolored green) and visualized under confocal microscopy (63X). (B) Relative Nucleoid Size (RNS) (larger axis; *inset*) of ~100-150 independent cells of various *Mtb* strains. For inducing WhiB4, 100 ng/ml of Atc was added to cultures of *whiB4- OE*. As a control, the same amount of Atc was added to other strains except HU. HU was overexpressed from an acetamide-inducible promoter as reported. Scale bar = 2 μm. Data shown are the average of two independent experiments done in triplicate. *** p ≤ 0.001 (as compared with *wt Mtb*).

We observed that the relatively expanded nucleoids in *wt Mtb* or *MtbΔwhiB4*, were compacted in cells overexpressing WhiB4 (*whiB4-OE*) (Fig. 3). The mean RNS value reduced by ~ 50% (0.125± 0.02 μm, p≤0.001) as compared to wt *Mtb* (0.261 ± 0.050 μm) or *MtbΔwhiB4*, (0.291± 0.05 μm). A similar level of compaction resulted from the overexpression of the well established mycobacterial NAP HU (45), which served as a positive control (Fig. 3). Native expression of WhiB4 in *MtbΔwhiB4*, (*whiB4-Comp*) resulted in nucleoid expansion to the wt *Mtb* level (Fig. 3). Furthermore, Atc-mediated over-expression of untagged WhiB4 condensed the mycobacterial nucleoid, ruling out the influence of the FLAG-tag on condensation mediated by WhiB4 (Fig. S6A). Further, our intracellular localization studies using indirect immuno-fluorescence and FLAG-specific antibody revealed that WhiB4 is exclusively associated with the DAPI-stained clumped nucleoids of *whiB4-OE* (Fig. 4). As an additional verification, we overexpressed the WhiB4-GFP fusion protein and confirmed that fluorescence remained associated with the compacted nucleoid (Fig. S6B). In contrast, over-expression of the cysteine mutant of WhiB4 (*whiB4- cys3-OE*) did not induce DNA condensation (Mean RNS = 0.255 μm ± 0.046, p ≤ 0.001 as compared to *whiB4-OE*), and the mutant protein was found scattered across the length of the cell (Fig. 4A and 4B). Expression of WhiB4 was maintained in the *whiB4-cys3-OE* strain, indicating that the loss of DNA condensation is due to disruption of the thiol-disulfide redox switch (Fig. 4C). Lastly, we studied WhiB4-induced nucleoid condensation by performing ultrastructure imaging of the mycobacterial nucleoid using Transmission Electron Microscopy (TEM; See Materials and Methods). Analysis of mid-logarithmic (log) phase cells overexpressing WhiB4 showed a highly condensed nucleoid, unlike the well spread out nucleoid in the wt *Mtb* cells (Fig. 5, compare C, D with A, B, respectively). The nucleoid morphology of the wt *Mtb* cells was as reported previously (46-48). On the contrary, the *MtbΔwhiB4*, mid-log phase cells possessed a highly unorganized and extensively spread out nucleoid that occupied the entire cell (Fig. 5E and F). The ultrastructural changes in *MtbΔwhiB4*, were rescued and the organization of the nucleoid was restored in *whiB4-Comp* (Fig. 5G and H). Altogether, using multiple analytical techniques, we confirmed that WhiB4 participate in mycobacterial nucleoid compaction in a redox-dependent manner.

**Figure 4:**
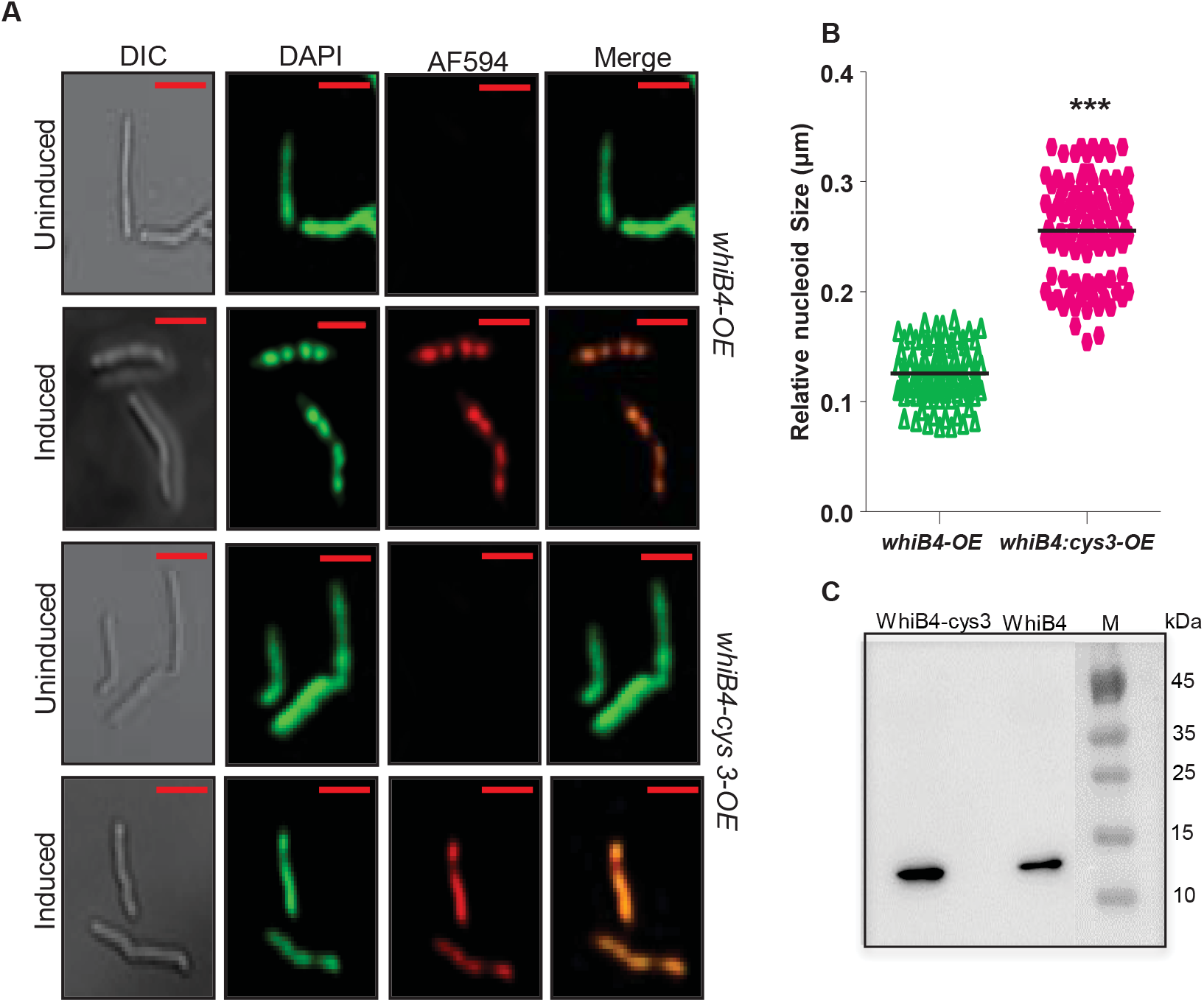
Cysteine residues of WhiB4 are important for its DNA condensation activity. **(A)** Confocal microscopic images (63X) of *MtbΔwhiB4*, strain over-expressing WhiB4- and WhiB4-cys3 FLAG tagged proteins. Nucleoids were stained with DAPI (pseudo colored green), wt WhiB4-FLAG and WhiB4-cys3-FLAG mutant were stained with the AF594 secondary antibody (red) against the anti-FLAG primary antibody, and co-localization was indicated by yellow in the merge panel. For inducing WhiB4, 100 ng/ml of Atc was added to cultures of *whiB4-OE and whiB4- cys3* strains. The uninduced (UI) control lacks immuno-fluorescence due to the absence of WhiB4 expression. The scale of images is 3 μm. **(B)** The RNS values of ~ 100-150 independent cells of various *Mtb* strains **(C)** 30 μg of cell free extract from either *whiB4-OE* or *whiB4-cys3-OE* strains was analyzed for the WhiB4 expression by immuno-blotting using the antibody against FLAG tag. Data shown are the average of two independent experiments done in triplicate *** p ≤ 0.001 (as compared with *wt whiB4-OE*).

**Figure 5:**
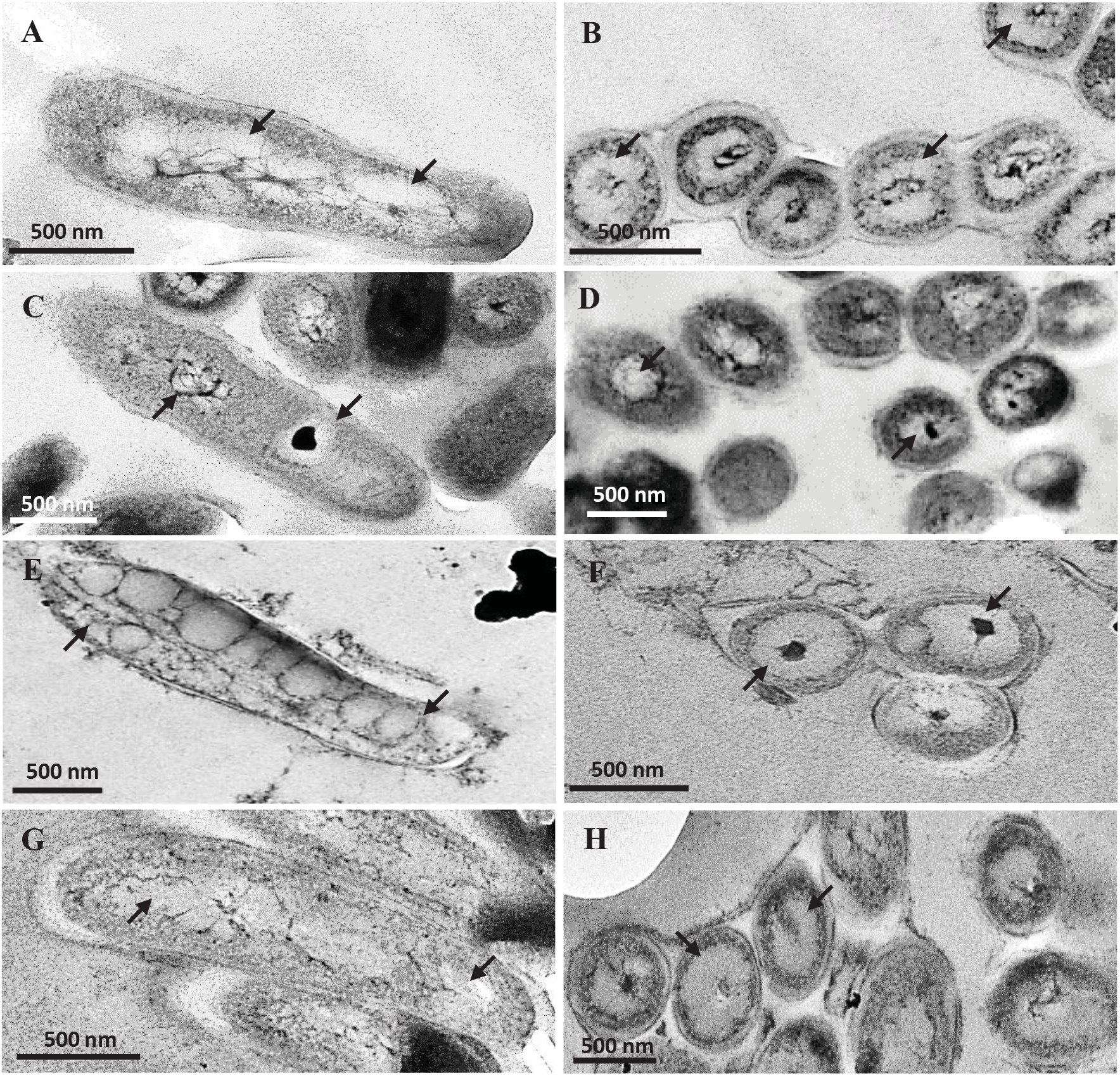
Transmission electron microscopic (TEM) analysis of *Mtb* nucleoids. TEM images of the longitudinal and transverse sections of: (**A, B**) wt *Mtb* from mid-log phase culture showing a well spread out nucleoid in the middle of the cell; **(C, D)** *whiB4-OE* cells showing highly condensed nucleoid in the middle of the cell; **(E, F)** *MtbΔwhiB4*, showing highly unorganized and well spread out nucleoid throughout the cell; **(G, H)** *whiB4-comp* showing well spread out nucleoid throughout the cell, like in (**A, B**). For inducing WhiB4, 100 ng/ml of Atc was added to cultures of *whiB4-OE*. As a control, the same amount of Atc was added to other strains. In each case, ~250-300 independent cells were visualized and representative images are shown. Arrowhead indicates nucleoid.

### WhiB4 regulates DNA condensation in response to oxidative stress

Having shown that WhiB4 expression modulates nucleoid condensation, we wanted to understand the influence of oxidative stress triggered by CHP on mycobacterial nucleoids and the role of WhiB4 in this outcome. We treated wt *Mtb*, *MtbΔwhiB4*, and *whiB4-OE* strains with 0.5 mM CHP and monitored redox potential, nucleoid condensation, and survival at various time points. To image the redox state of *Mtb*, we measured the redox potential of it’s most abundant cytoplasmic thiol (mycothiol; MSH) using Mrx1-roGFP2. The biosensor shows an increase in fluorescence excitation ratio at 405/488 nm upon oxidative stress, whereas a ratiometric decrease is associated with reductive stress (49). The ratiometric changes (405/488 nm) in the fluorescence of the biosensor can be fitted to the modified Nernst equation to precisely determine the millivolt (mV) changes in the redox potential of mycothiol (*E_MSH_*) (49).

Both wt *Mtb* and *MtbΔwhiB4*, displayed a steady-state *E_MSH_* of ~ -276 mV. However, overexpression of *whiB4* induces an oxidative shift in *E_MSH_* of *whiB4-OE* (~-250 mV), consistent with the antioxidant repressor function of WhiB4. Treatment with CHP induces a significant oxidative shift in *E_MSH_* of wt *Mtb* (-220 to -225 mV) at 6 h and 24 h post-treatment (Fig. 6A). Importantly, the nucleoids of wt *Mtb* underwent a noteworthy condensation at 6 h (mean RNS: 0.14 μm ± 0.02) and 24 h (mean RNS: 0.10 μm ± 0.01) post-treatment (Fig. 6A). At 48 h post CHP-treatment, a majority of the cells showed loss of DAPI fluorescence and appeared rounded/irregular, thus precluding any measurements of nucleoid length. These morphological alterations are indicative of DNA fragmentation and killing, which we confirmed by CFU analysis (Fig. 6B) and SYTO9-propidium iodide staining (Fig. S7). Using indirect immunofluorescence assays and the WhiB4-GFP fusion, we confirmed that WhiB4 expressed from its own promoter or overexpressed remained associated with the condensed nucleoid upon CHP treatment at 6 h and 24 h post-treatment (Fig. S8). Notably, the skew towards condensed DNA and oxidative *E_MSH_* was activated at a time point where toxicity was marginal (*i.e.* 6 h), indicating that genome hypercompaction likely precedes oxidative stress-induced killing of *Mtb*.

**Figure 6:**
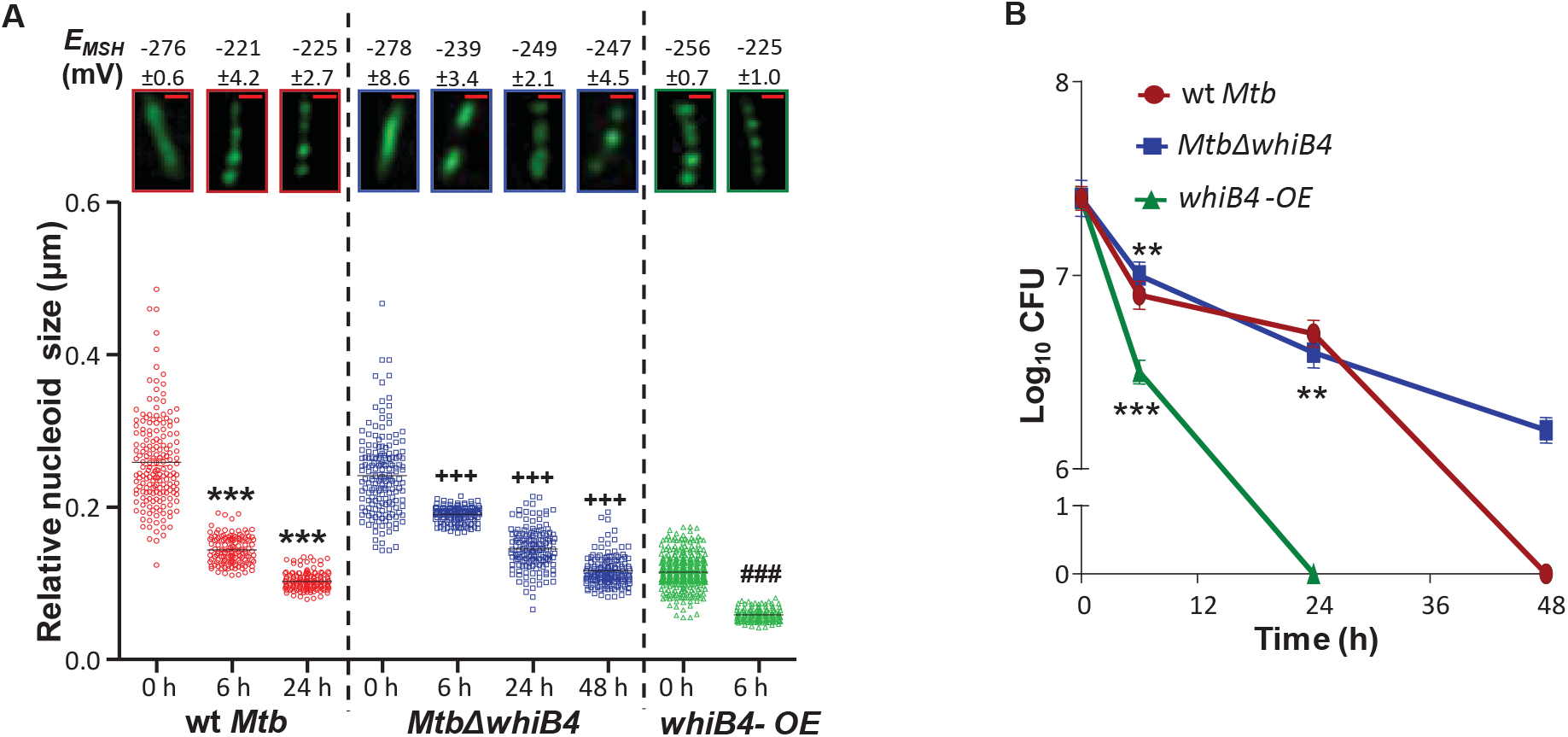
Oxidative stress leads to DNA condensation, skewed redox homeostasis and bacterial killing in a WhiB4-dependent manner. wt *Mtb, MtbΔwhiB4*, and *whiB4-OE* were exposed to 0.5 mM CHP for 6, 24 and 48 h. Nucleoids were stained with DAPI (pseudo colored green) and visualized by confocal microscopy (63X). The scale of images is 1μm. For inducing WhiB4, 100 ng/ml of Atc was added to cultures of *whiB4-OE*. As a control, the same amount of Atc was added to other strains. The relative RNS of ~ 100-150 independent cells was measured and represented as scatter plots. Each dot represents one nucleoid. The line depicts mean of the population at each time point. *P values*: * = as compared to *wt Mtb,* + = as compared to *MtbΔwhiB4*, # = as compared to *whiB4-OE* at 0 h (\*\*\**P* or ^+++^*P* or ^###^*P* ≤ 0.001). Intramycobacterial *E_MSH_* of various *Mtb* strains at each time point was measured using flowcytometer and shown along with a corresponding image of a representative DAPI stained cell visualized by confocal microscopy under similar conditions. **(B and D)** Plots showing the survival of *Mtb* strains in response to CHP stress as assessed by enumerating CFUs. Data shown is the average of experiments performed in triplicates. Error bars represent SD from the mean.

In contrast to wt *Mtb*, nucleoids of *MtbΔwhiB4*, underwent a slower transition towards a highly condensed state, which corresponds to a moderate shift towards oxidative *E_MSH_* (*E_MSH_*: -247 to -239 mV) and a relatively much slower killing by CHP (Fig. 6A and 6B). The fact that *MtbΔwhiB4*, exhibits nucleoid hypercompaction, albeit delayed, indicates the role of additional factors in nucleoid compaction during oxidative stress. Not unexpectedly, 0.5 mM CHP treatment for 6 h was sufficient to induce substantial DNA condensation (0.06 μm ± 0.01), overwhelm redox balance (*E_MSH_*: -225 mV ± 0.98) and induce killing in the *whiB4-OE* strain (Fig. 6A and 6B). Measurements of nucleoid length of the *whiB4-OE* strain at 24 and 48 h post-CHP treatment could not be performed due to massive cell death (Fig. 6B). It can be argued that the high concentration of CHP (0.5 mM) can adversely affect *Mtb’s* physiology to influence genome topology. To address this issue, we reassessed nucleoid condensation, intrabacterial *E_MSH_*, and survival upon exposure to non-toxic concentration of CHP (0.1 mM) over time. A relatively moderate effect of 0.1 mM CHP was observed on nucleoid condensation*, E_MSH_*, and survival. However, the relative differences in nucleoid condensation*, E_MSH_*, and survival between wt *Mtb*, *MtbΔwhiB4*, and *whiB4-OE* followed the order obtained with 0.5 mM CHP (SI note IV; Supplementary Fig. S9).

Since our results indicate a dynamic relationship between oxidative stress and genome condensation, we examined the state of nucleoids in a *Mtb* strain completely devoid of MSH antioxidant (*MtbΔwhiB4*,) and in a *mshA* complemented strain (*mshA-comp*) (50). Similar to the *whiB4-OE* strain, *MtbΔwhiB4*, grows normally under aerobic growing conditions but shows oxidative *E_MSH_* and acute sensitivity towards oxidative stress (51). In agreement to findings with *whiB4-OE*, nucleoids of *MtbΔwhiB4*, cells showed hypercondensation (mean RNS: 0.11 μm ± 0.04) as compared to the *mshA-comp* cells (mean RNS: 0.25 μm ± 0.02) (Fig. S10) under normal growing conditions. These results reinforce a functional association between DNA condensation and tolerance to oxidative stress in *Mtb*. In summary, our data indicate that deletion of WhiB4 reduces, and overexpression potentiates, the adverse impact of DNA condensation on oxidative stress survival.

### ChIP-Seq demonstrates a nearly-uniform association of WhiB4 with *Mtb* chromosome

To determine genome-wide distribution of oxidized apo-WhiB4, we performed ChIP-Seq. Native expression of WhiB4 predominantly generates non-DNA binding forms (holo-WhiB4 and/or reduced apo-WhiB4) *in vivo* (Fig. S2B), consistent with a marginal effect of WhiB4 on gene expression under normal growth conditions (22). Secondly, while we can oxidize WhiB4 by CHP treatment in *Mtb*, oxidized apo-WhiB4 also functions as an autorepressor and therefore can present challenges in identifying weak or transient binding sites. Therefore, we utilized ectopically expressed FLAG-tagged WhiB4 (*whiB4-OE*), which bypasses autoregulatory loop and consistently generates oxidized apo-WhiB4, to perform ChIP-Seq. We considered the possibility that ectopic expression might result in nonphysiological DNA binding. However, our data showed that over-expression of WhiB4 using 100 ng of Atc produced nearly physiological concentrations of oxidized apo-WhiB4 molecules/cell (comparable to CHP treated *Mtb*) (Fig. S2 and S5). Furthermore, using qRT-PCR we confirmed that Atc-mediated overexpression of FLAG-tagged or untagged WhiB4 influences the expression of only WhiB4-specific genes (S2 table) and affects pathogen’s survival specifically under oxidative stress *in vitro*, *ex vivo*, and *in vivo* (*SI Note III* and Fig. S11). In addition, a recent study showed notable overexpression of *whiB4* during starvation and upon hypoxia (52, 53). Lastly, recent studies have demonstrated remarkable consistency between the DNA binding profile of several ectopically overexpressed FLAG-tagged transcription factors and ChIP-Seq studies performed under native conditions in *Mtb* (54). Thus, while we cannot exclude the possibility that overexpression could lead to nonphysiological DNA binding, we conclude that overexpression did produce physiologically-relevant redox-form(s) of WhiB4 necessary for DNA binding in *Mtb*.

We induced the expression of WhiB4 using 100 ng of Atc for 16 h, harvested chromatin samples for ChIP-Seq using anti-FLAG antibody conjugated to magnetic beads, and sequenced the cross-linked DNA using the Illumina Genome Analyzer system (Materials & Methods). As a negative control, we sequenced the input chromatin sample prior to immunoprecipitation (IP) with the anti-FLAG antibody. In parallel, we performed the analysis of a published genome-wide DNA binding study that was conducted using a sequence-specific transcription factor, CRP, in *Mtb* (55). For each sample, we examined read count distribution by measuring the number of reads mapped to each base on the *Mtb* chromosome and then plotted the densities of read count distributions. The data for two independent samples of WhiB4 showed a weakly skewed distribution to the right, which is largely similar to that obtained from input experiment (Fig. 7A, 7B, 7C, and 7D). In agreement to this, the read counts obtained for WhiB4 showed a more significant correlation with the input (*ρ* = 0.45 and 0.379), whereas read-counts for CRP were noticeably skewed towards the right and showed a weak correlation with the input control (*ρ* = 0.245, Fig. 7E and 7F). Although the binding profile from the WhiB4 ChIP-seq experiment showed a strong resemblance to that from the input, we think that the data is representative of DNA binding profiles of WhiB4. For example, the read count profiles for each WhiB4 replicate are more correlated with each other (*ρ* = 0.81; Fig. 8A) than with the input (*ρ* = 0.45 or 0.379). Secondly, although weak, there is a clear right-sided skew in case of the ChIP signal as compared to the input. Finally, ChIP experiments for WhiB4 were consistently successful and obtained high concentrations of DNA (8.54 ng/μl and 8.52 ng/μl) as compared to mock-IP (2.72 ng/μl and 3.12 ng/μl) or the WhiB4-cys3 mutant (3.31 ng/μl and 2.98 ng/μl), which generally provides inadequate concentrations of DNA for a sequencing reaction. Altogether, the data support our *in vitro* findings suggesting a more uniform binding of WhiB4 to the chromosome of *Mtb*.

**Figure 7:**
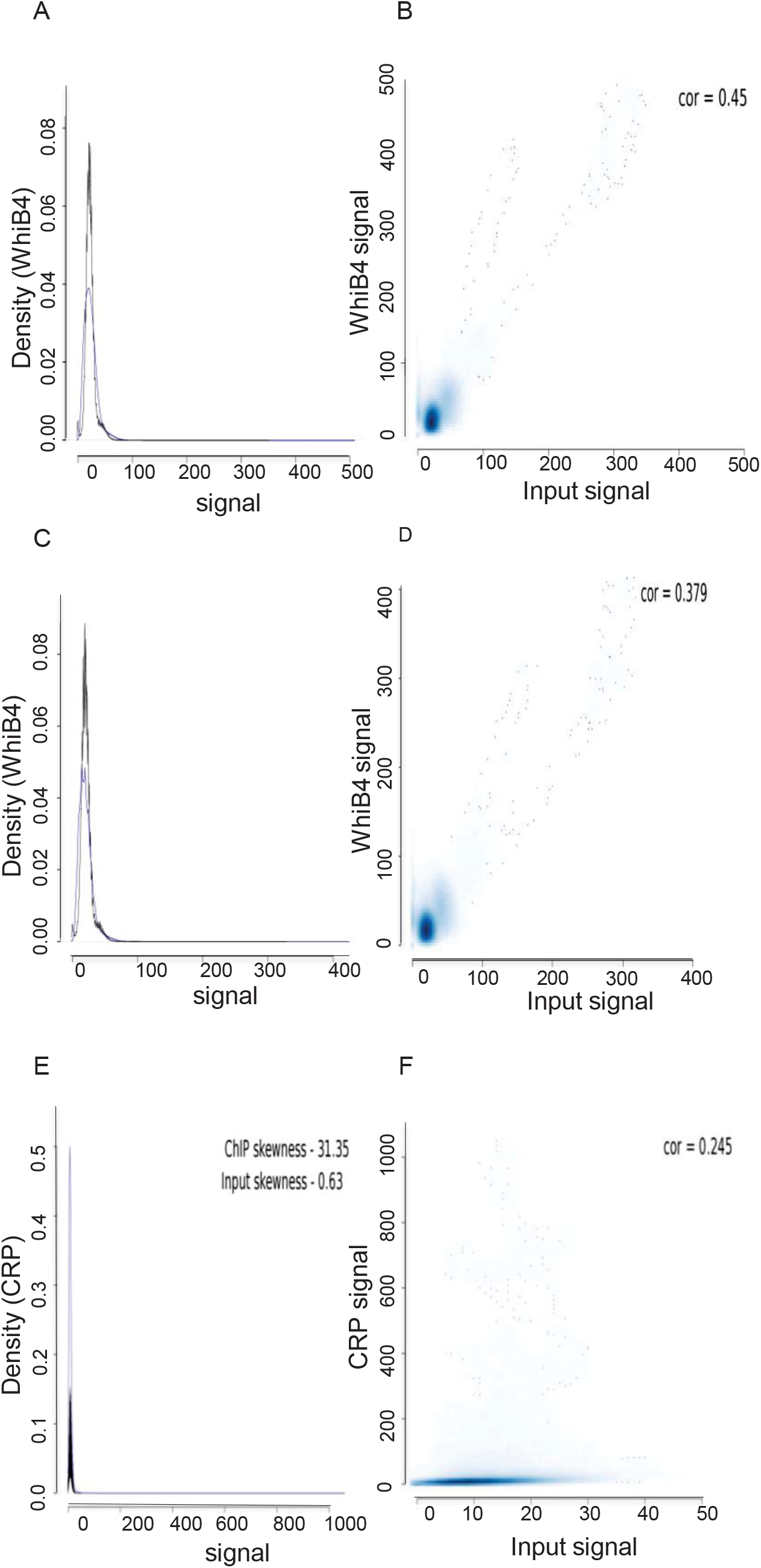
WhiB4 exhibits largely uniform binding to the *Mtb* chromosome. The left panel shows density distribution of the read counts (*x*-axis) of (A) WhiB4 replicate 1, (B) WhiB4 replicate 2, and (C) CRP ChIP-Seq (in blue) and the Input (in black). Compared to CRP, the distribution of the WhiB4 biological replicates distribution does not have the characteristic heavy right end tail. This is also evident from the skewness measured for each distribution. Positive values depict right-handed distribution as in the case of CRP. The right panel shows the scatterplot along with the correlation (at single base resolution) in read counts between the (B) WhiB4 replicate 1, (D) WhiB4 replicate 2, and (F) CRP ChIP-Seq and their respective inputs.

We next characterized binding by obtaining a measure of the WhiB4 occupancy for each gene in the genome using the method used to quantify nucleosomal occupancy in eukaryotes and binding of a NAP HU in prokaryotes ((56) and (57); see Materials and Methods). This analysis revealed that WhiB4 occupancy positively correlates with the G+C content of the bound DNA (Fig. 8B), which is in agreement with our earlier work on *in vitro* specificity of WhiB4 interaction with a specific locus (*ahpCD*) (22). As expected, a highly GC-rich motif was identified by MEME for WhiB4 binding (Fig. 8C, E-value: - 6.4×10^-103^). For two independent WhiB4 samples, approximately 700 regions of enriched signals with a remarkable (>90%) overlap between the binding regions were obtained (S3 Table). Furthermore, genomic regions bound by WhiB4 are longer than the stretches of DNA bound by CRP (Fig. 8D). Approximately, 7-10% of the *Mtb* genome encodes PE (99) and PPE (69) proteins (58, 59). Of the 90 PE proteins, 63 were encoded by GC-rich genes and they belong to the PE-PGRS sub-class. Expectedly, WhiB4 bound overwhelmingly to PE-PGRS genes (55 of 63), but marginal to PE/PPE genes that do not belong to the PGRS sub-class. This result is consistent with the observed preference of WhiB4 for GC rich regions (Fig. 9 and S3A Table) in *Mtb* and WhiB4-mediated regulation of PE-PGRS expression in *Mycobacterium marinum* (60). Additionally, genes involved in respiration (*e.g.,* cytochrome BD oxidase, NADH dehydrogenase), redox homeostasis such as mycothiol biosynthesis (*mshA, mshB, mshD, mrx1*), thioredoxin pathway (*trxB2, thiX*), NAD metabolism (*nadA, -B, -C, nudC*), and Fe/Fe-S hemostasis (*sufR, sufB, bfrB, mbtB*) were also bound by WhiB4 (S3A Table). Interestingly, WhiB4 binds to several transcription factors, sigma factors, and NAPs (*lsr2, espR, IHF*) (S3A Table). To validate the WhiB4 binding peaks, we performed ChIP followed by quantitative PCR (ChIP-qPCR) on 9 target DNA sequences (5 within intergenic regions, 3 within ORFs and 1 non-peak region). Consistent with the ChIP data, all of the selected WhiB4-binding regions showed enrichment (S4 Table). As a negative control, we over-expressed the Flag-tagged WhiB4-Cys3 variant and performed ChIP-qPCR of the WhiB4 targets on immuno-precipitated genomic DNA as described earlier. ChIP-qPCR of WhiB4-specific regions showed no enrichment in the case of WhiB4-Cys3 (S4 Table). Thus, the results indicated that all peaks were likely to be genuine WhiB4-targets.

**Figure 8:**
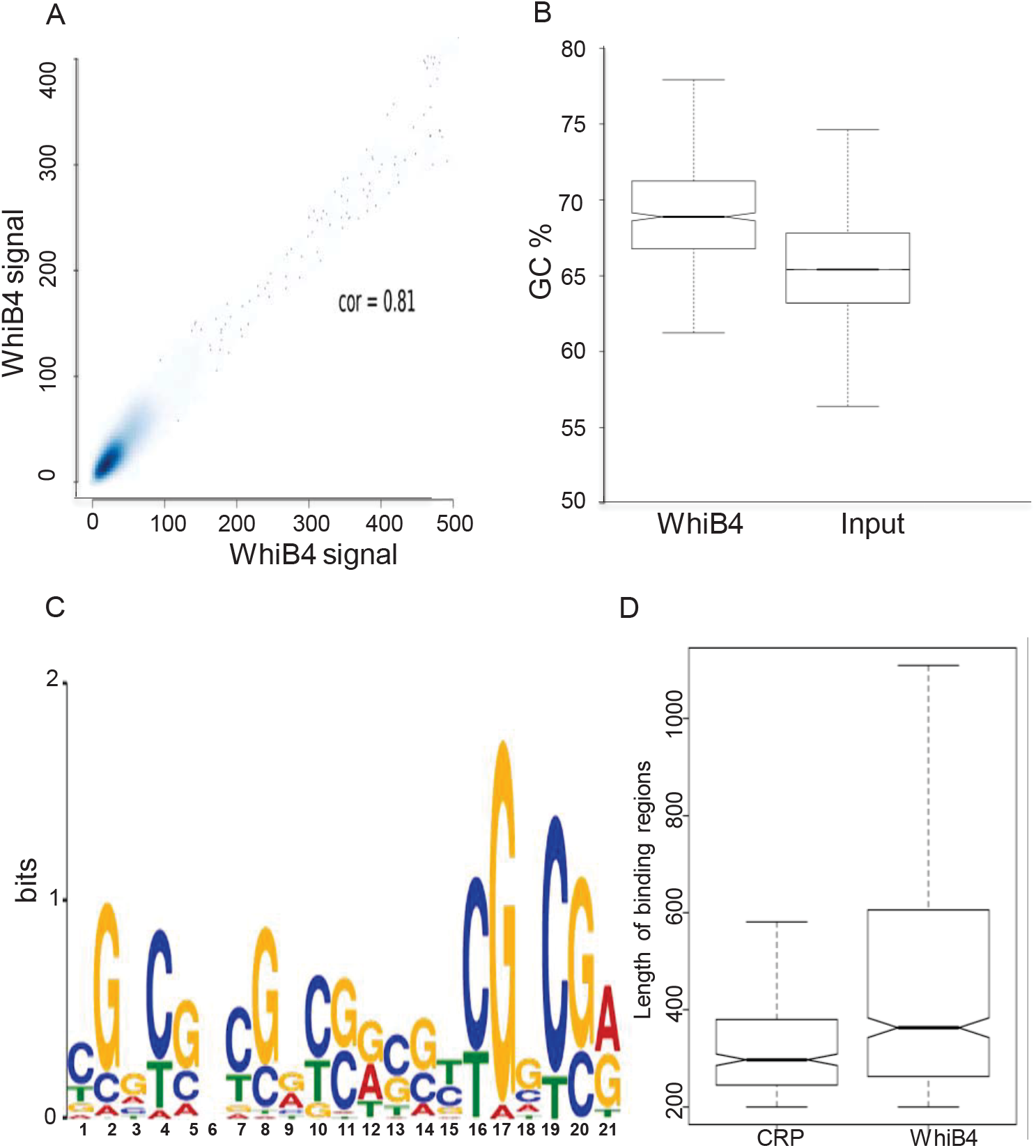
Length and AT/GC preference of WhiB4 binding. (A) Scatterplot of the correlation between WhiB4 biological replicates. (B) Boxplot showing GC% in WhiB4 peaks and comparison with the randomized input regions. The enrichment in the binding regions is statistically significant (P < 0.01). (C) GC rich Motif enriched in the WhiB4 binding regions as determined by MEME-ChIP. (D) Length distribution of WhiB4 binding regions in the boxplot and comparison with the available CRP dataset. On the y-axis is the number in base pairs.

**Figure 9:**
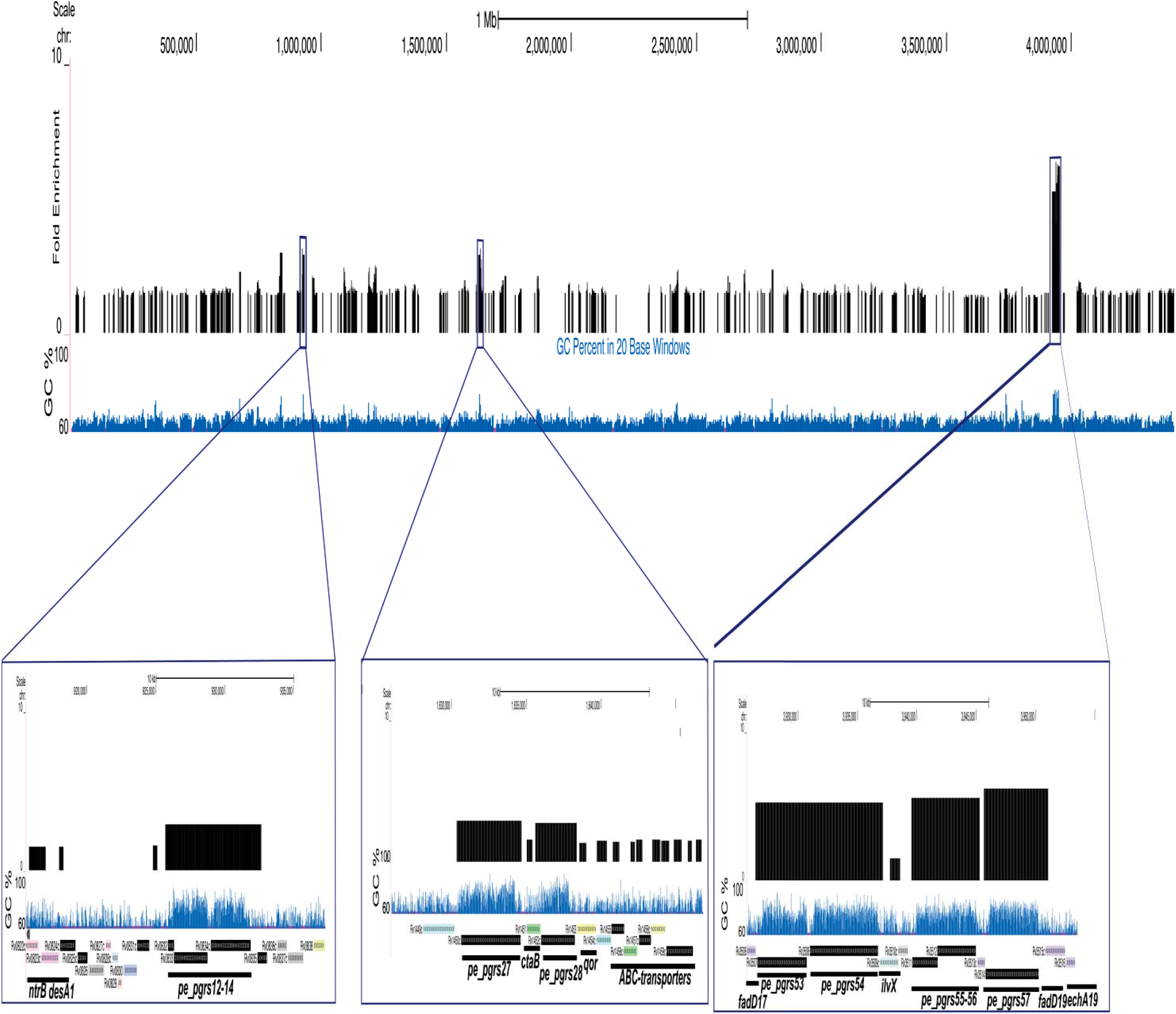
Genome wide mapping of WhiB4 binding sites. Input normalized WhiB4 binding (in black) on the chromosome as visualized on the UCSC genome browser and the corresponding GC%. On the y-axis, the fold enrichment of the bound regions, except for few regions remains at low level. High fold enrichment was found for genomic regions rich in GC content as seen in the zoomed in examples.

Finally, we investigated whether genes bound by WhiB4 showed differential expression in *Mtb* during CHP stress. By comparing microarray of CHP-treated *Mtb* (over untreated *Mtb*) with the ChIP-seq data, we found that expression was lower, albeit marginally, for genes bound by WhiB4 than those that are not (Fig. S12, Supplementary table S3B; Wilcoxon test, *P* < 2.2 x10^-16^). While these results showed an association between WhiB4 binding and repression of corresponding genes, they do not establish causality. To understand this, we assessed the genome wide binding data and the microarray data for CHP-treated *Mtb∆whiB4* (over CHP-treated *Mtb*). In this analysis, the large number of genes differentially expressed in CHP-treated *Mtb∆whiB4* accounts for ~ 30 % of the WhiB4-bound genes (Fig. S12, Supplementary table S3C). These findings suggest that the deletion of *whiB4* indirectly influences the expression of a large number of genes (~ 70%) affected by CHP. This insufficiency could be due to counterbalancing effect of other transcription factors/NAPs on the expression of these genes. Similar to our conclusions, studies on Fis and Crp in *E. coli* have found little association between Fis/Crp DNA binding and differential gene expression in *∆fis/∆crp* strains (61-63). It was proposed that the primary function of *E. coli* CRP and FIS proteins is to sculpt the chromosome, while their role in regulating transcription is likely to be circumstantial (61-63). A similar explanation may be suited for WhiB4 as well.

## Discussion

In this study, we identified a link between DNA condensation, WhiB4, and oxidative stress response in *Mtb*. WhiB proteins are proposed to be Fe-S cluster-containing transcription factors; however, the exact molecular mechanism of action remained poorly understood. Studies revealed that some of the WhiB family members might regulate gene expression by binding to promoter sequences (20, 22, 64). Furthermore, ChIP-Seq analysis of WhiB in *Streptomyces coelicolor* (*Sco*) seems to indicate DNA binding wherein a specific regulatory effect may be achieved by its association with a transcription factor WhiA (65). Using various experimental approaches, we confirmed that WhiB4 condenses DNA by binding more uniformly to the *Mtb* chromosome, with a notable preference for GC-rich DNA. Our expression and ChIP-Seq data showed that the WhiB4 regulon is composed of genes involved in redox homeostasis, CCM, respiration, PG metabolism, PE-PGRS, and Esx-1 secretory virulence factors. Over-expression of WhiB4 induced hypercondensation of nucleoids *in vivo*, indicating that the protein, in sufficient quantity, can function independently in regulating DNA compaction. Notably, over-expression of NAPs (*e.g*. Dps) does not always result in nucleoid condensation owing to the negative influence of other NAPs (*e.g*. Fis) (30). While overexpression of WhiB4 can condense nucleoid *in vivo*, how it contributes to DNA condensation in conjunction with other mycobacterial NAPs remains to be determined. Moreover, a large majority of WhiB4-binding events are inconsequential from the transcriptional perspective, suggesting that WhiB4-binding does not necessarily affect transcription locally, but may serve as focal points to organize the genome into domains and thereby influence transcription indirectly. Association of WhiB4 with genomic regions encoding Lsr2, EspR, and IHF does indicate the interplay between different NAPs to alter chromosome structure and organization, thereby influencing patterns of gene expression in response to oxidative stress.

Several NAPs are known to play important role in influencing nucleoid condensation under specific stress conditions such as iron starvation, hypoxia, and acidic pH (9). However, the importance of DNA condensation and NAPs in controlling oxidative stress response remains controversial. For example, while *S. aureus* resists H_2_O_2_ by rapidly condensing DNA through a Dps homolog (MrgA) (12), oxidative stress simply did not lead to DNA condensation in other bacteria including *Sco*, *Dickeya dadantii*, and *E. coli* (30, 66, 67). Interestingly, a recent study elegantly showed that MrgA-mediated protection of *S. aureus* from oxidative stress is mainly due to its ferroxidase activity rather than its ability to shield DNA (68). However, the role of Lsr2 in protecting *Mtb* from oxidative damage by shielding DNA is controversial (11, 69). Therefore, it is unclear as of now how mycobacteria remodel their DNA condensation during oxidative stress. To begin understanding this, we demonstrated that the loss of WhiB4 delays DNA condensation, maintains redox homeostasis, and protects the cells from oxidative stress, whereas its over-expression reversed these phenotypes. Our data indicate that the detrimental effect of WhiB4 overexpression is likely to be due to the repression of antioxidant machinery and the diminished capacity to counteract cytoplasmic redox stress. It is also likely that hyper-condensation of the nucleoid inhibits other metabolic processes such as DNA replication/repair, transcription, and translation to exert efficient killing. We conclude that resistance to oxidative stress in *Mtb* is unlikely to be mediated by nucleoid hypercompaction. In this context, it has been suggested that certain bacteria (*e.g.,* gamma-proteobacteria) have evolved genetic mechanisms (*e.g.* Fis, TopA, and GyrA) to block DNA condensation and promote the expression of OxyR-dependent antioxidant genes as a major mechanism to guard genomic DNA against oxidative stress (30). Moreover, similar to *Mtb*, a higher degree of oxidative stress and killing were associated with nucleoid hypercondensation in *E. coli* (15, 70). Although, unlike *Mtb* where WhiB4 overexpression is sufficient to induce both DNA condensation and killing under oxidative stress, a similar consequence of oxidative challenge seems to be mediated through a combined action of multiple OxyR-regulated NAPs (*e.g*, H-NS, Hup, Him, MukB, and Dps) in *E.coli* (15). In the absence of OxyR, Fis and Dps activities in *Mtb*, WhiB4, with its redox-active cysteines can be an important regulator of both nucleoid condensation and expression of oxidative stress responsive genes in *Mtb*.

We propose that under aerobic growing conditions, air oxidation of holo-WhiB4 to oxidized apo-WhiB4 activates DNA binding and repressor function to preclude the unnecessary expression of oxidative stress responsive pathways. A gradual increase in oxidative stress on the one hand increases generation of oxidized apo-WhiB4 oligomers while on the other hand down-regulates WhiB4 expression. Both of these activities ensure levels of intracellular oxidized apo-WhiB4 oligomers required to induce appropriate degree of nucleoid condensation and calibrated activation of antioxidants/stress pathways. The steady increase in antioxidants such as MSH and Fe-S biogenesis machinery (Suf operon) in response to sustained oxidative stress can create a regulatory feedback loop by reducing oxidized thiols of apo-WhiB4 to regenerate monomeric reduced apo-WhiB4 and/or holo-WhiB4. This would allow WhiB4 to loose chromosomal binding and completely derepress antioxidants, maintaining both redox and topological balance (Fig. 10). Besides this, other factors (*e.g.,* NAPs, sigma factors *etc*) which directly or indirectly cooperate with WhiB4 could play a role in maintaining topological homeostasis and adaptation to oxidative stress. The fact that *Mtb∆whiB4* showed higher resistance to oxidative stress and *whiB4-OE* displayed hypersensitivity as compared to wt *Mtb* indicates that a gradual decrease in *whiB4* expression upon oxidative stress is a cellular decision to induce the oxidative stress response and reduce DNA hypercondensation.

**Figure 10:**
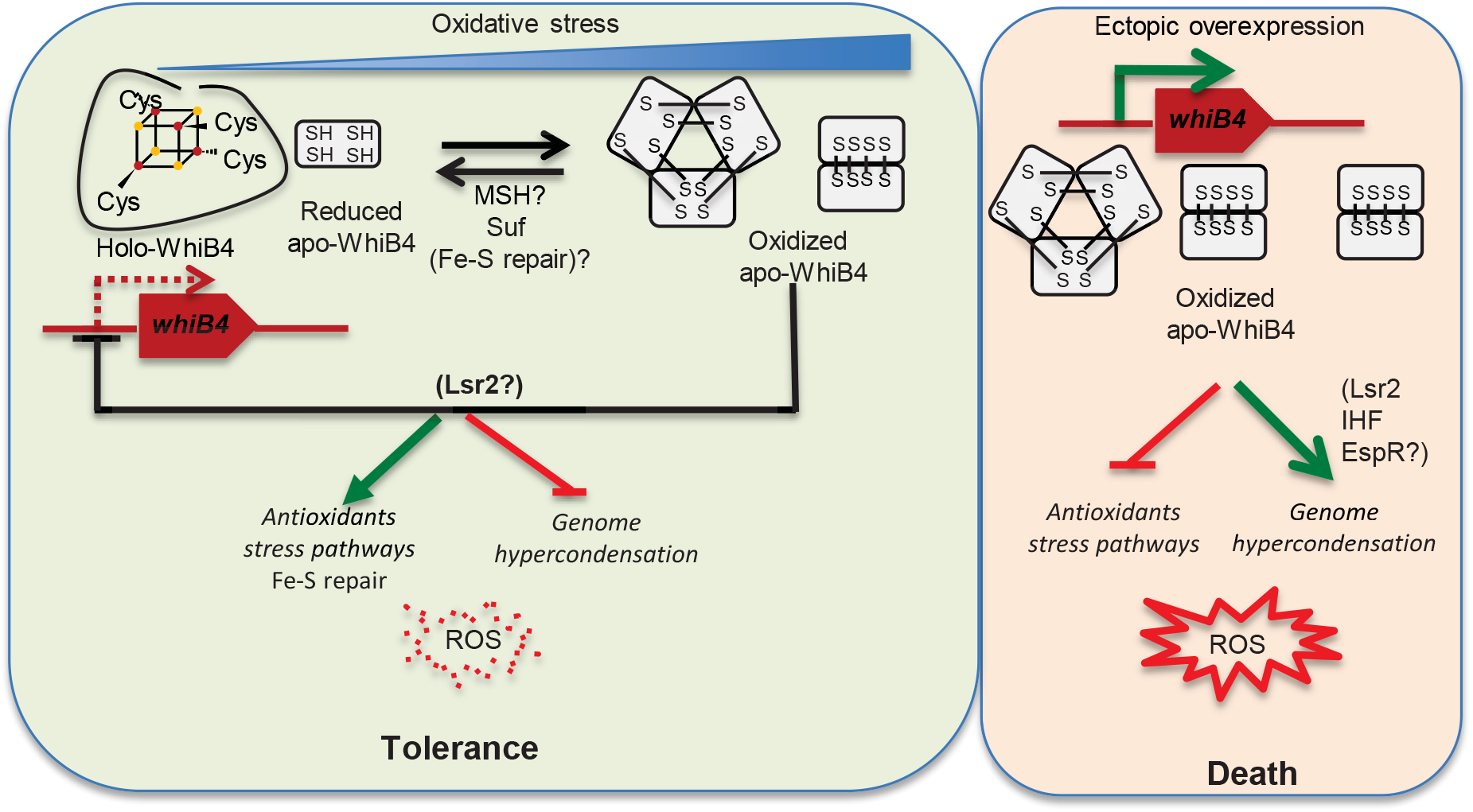
Proposed model of WhiB4 function in oxidative stress response. Under mild oxidative conditions, a minor fraction of oxidized apo-WhiB4 generated via oxidation of holo-WhiB4 activates DNA binding and repressor function to preclude unnecessary overexpression of oxidative stress responsive pathways. Further elevation in oxidative stress levels result in higher thiol-linked apo-WhiB4 oligomers, which would have further repressed antioxidants expression and induced genome hypercondensation. To circumvent this, *Mtb* downregulates expression of WhiB4 (via autorepression and/or Lsr2) to control the levels of oxidized apo-WhiB4, which ensures appropriated degree of nucleoid condensation and antioxidants expression. A gradual increase in antioxidants such as MSH and the Fe-S repair pathways (Suf operon) can create a regulatory feedback loop, which reduces oxidized thiols of apo-WhiB4 to eventually regenerate monomeric reduced apo-WhiB4/holo-WhiB4. All of this ultimately results in complete derepression of antioxidants to mitigate oxidative stress and to also restore topological homeostasis. Ectopic over-expression of WhiB4 by Atc leads to sustained accumulation of oxidized apo-WhiB4 without any feedback control. This eventually represses antioxidants/stress pathways and induces long lasting nucleoid hypercondensation. Both of these activities of WhiB4 adversely affect mycobacterial ability to tolerate oxidative stress leading to enhanced susceptibility of WhiB4-OE strain towards CHP.

Recently, starvation-induced dynamic changes in the mycobacterial nucleoid content and structure were shown to trigger a development program resulting in the formation of a small-cell morphotype with hypercondensed nucleoid, elevated antibiotic tolerance, and persistence capabilities (71). More importantly, RNA-seq analysis of nutritionally-deficient cells containing condensed nucleoids has shown a remarkable over-expression of *whiB4* (52). Therefore, WhiB4-directed nucleoid hypercondensation might mediate development and differentiation in *Mtb* under dormancy-inducing conditions to facilitate long term persistence (work currently under progress). In conclusion, the combined results indicate that WhiB4 is a candidate for a dual-function NAP that could integrate environmental signals with DNA conformation and transcription. The focus of the present investigations has been mainly on the nucleoid condensing function of WhiB4 and how it can modulate oxidative stress response in *Mtb*. As far as we are aware, this is the first example of a redox-dependent NAP in bacteria.

## Materials and Methods

### Bacterial strains and growth conditions

Various *Mtb* strains and primers used in this study were listed in S5 Table and were cultured as described previously (22). *E. coli* cultures were grown in Luria-Bertani (LB) medium (BD Biosciences). When required, the culture medium was supplemented with hygromycin (50 μg/ml for mycobacteria, 150 μg/ml for *E.coli*), kanamycin (25 μg/ml for mycobacteria and 50 μg/ml for *E.coli*), and ampicillin (100 μg/ml). For cumene hydroperoxide (CHP, Sigma Aldrich) stress, strains were grown to an exponential phase and exposed to different CHP concentrations. Survival was monitored by enumerating colony forming units (CFU) at 0, 6, 24, and 48 h post-treatment. To examine the influence of WhiB4 over-expression on stress tolerance, *whiB4-OE* strain was grown aerobically till O.D._600_ of 0.3, induced with 200 ng/ml Anhydro Tetracycline (Atc, Cayman Chemicals) for 24 h at 37°C, and exposed to (i) normal aerobic environment, (ii) acidic stress (pH 4.5), and (iii) heat stress (42°C). The growth kinetics was monitored over time by measuring absorbance at 600 nm.

### Atomic force microscopy (AFM)

The oxidized apo-WhiB4 was incubated with 7ng/ μl of supercoiled or relaxed forms of plasmid DNA (pEGFP-C1) in a concentration range from 1:2.5 to 1:20 (DNA: WhiB4; w/w) at RT and 10μl of this solution was loaded onto freshly cleaved mica surface. A similar procedure was followed for Cys3-WhiB4 following incubation with DNA at RT. The 10μl of protein-DNA mix was allowed to spread spontaneously and incubated for 1 min to allow the molecules to adhere on the mica surface. The unbound material was washed with deionised water and the bound surface allowed to air dry. Imaging was carried out using the 5500 scanning probe microscope (Agilent Technologies, Inc.) and the PicoView software. Images were obtained in tapping mode in the air with 225-mm-long silicon cantilevers (Agilent Technologies) that have a resonance frequency of 75 kHz and a force constant of 2.8 Newton/m. Scan speed used was 1 line/s. Minimum image processing (first order flattening and brightness contrast) was employed. Image analysis was performed using Pico Image software v1.4.4.

### Transmission electron microscopy

Cells were processed for transmission electron microscopy (TEM), as described previously (47, 72). Cells were prefixed with 1% (w/v) osmium tetroxide buffered in 0.15 M cacodylate buffer (pH-7.2) (Sigma) for 1 hr at room temperature. The prefixed cells were then washed once with the same buffer and post fixed for 2 hrs at room temperature in 0.15 M sodium cacodylate (Sigma) buffer containing 2% (w/v) tannic acid (Sigma) and 2% (v/v) glutaraldehyde (Sigma). Subsequently, the cells were subjected to washing with the same buffer, and were refixed in 1% (w/v) osmium tetroxide overnight at 4ºC. Cells were dehydrated in a graded series of ethanol solutions (Merck Millipore) ranging from 20% to 100% with an incubation period of 10 min at each step and finally embedded in LR White resin (Electron Microscopy Sciences) overnight. The embedded samples were then cut with a glass knife using an ultramicrotome by maintaining the section thickness at 70 nm. The sections were stained with 0.5% uranyl acetate (Sigma) and 0.04% lead citrate (Fluka), and observed using FEI Tecnai™ G2 Spirit electron microscope at 120 kV.

### E_MSH_ measurements

Measurements of intrabacterial *E_MSH_* during growth *in vitro*, upon exposure to CHP stress, were performed as described previously (49).

### Miscellaneous procedures

Additional materials and methods are provided in the supporting information.

### Ethics Statement

This study was carried out in strict accordance with the guidelines provided by the Committee for the Purpose of Control and Supervision of Experiments on Animals (CPCSEA), Government of India. The protocol was approved by the Committee of the International Centre for Genetic Engineering and Biotechnology, New Delhi, India (Approval number: ICGEB/AH/2011/2/IMM-26). All efforts were made to minimize the suffering.

## Acknowledgements

We are thankful to the University of Delhi South Campus MicroArray Centre (UDSCMAC), New Delhi, for conducting the microarray experiments. We thank IISc and ICGEB for providing BSL3 facilities. We are also grateful to the Imaging Facility at University of Delhi, South Campus, New Delhi, for all the confocal microscopy experiments and analysis. We are thankful to Dr. N. Ganesh, Department of Biochemistry, IISc, Bangalore, Dr. Anjana Badrinarayanan, Massachusetts Institute of Technology, USA, and Dr. S S Shivakumara, IBAB, Bangalore for their comments and feedback.

## Supporting Information

### SI Note I

#### *In vivo* redox state of WhiB4 thiols upon CHP stress

To understand the effect of oxidative stress on WhiB4, we quantified the redox and/or oligomeric state of WhiB4 in response to CHP stress *in vivo*. To accomplish this, we inserted His-tag at the N-terminus of WhiB4 and His-WhiB4 was cloned in an *Escherichia coli-*mycobacterial integrative vector, pCV125, and expressed it in *MtbΔwhiB4* from its native promoter. Since cysteine thiols are sensitive to oxidation, we pretreated the cells with a cell-permeable thiol-alkylating agent, N-ethylmaleimide (NEM), and prepared cell-free extracts of mid-exponential phase grown cultures for non-reducing SDS-PAGE and western blot analysis using anti-His antibody. NEM treatment effectively clamps the intracellular redox state of thiols, thereby preventing oxidation artifacts induced during cell-free extract preparation.

*In vitro* studies performed earlier have shown that oxidation of holo-WhiB4 resulted in the loss of Fe-S cluster and generation of apo-WhiB4. The apo-WhiB4 further oxidizes to form SDS-resistant dimers and trimers due to the formation of intermolecular disulfide bonds between its cysteine thiols (1). However, the redox and oligomeric state of WhiB4 in *Mtb* have never been characterized *in vivo*. Similar to the *in vitro* results, western blot analysis of cell-free extract yielded three bands of (~ 14 kDa, ~ 28 kDa, and ~ 42 kDa) that corresponded to the size of a full-length His-tagged apo-WhiB4 monomer, dimer and trimer, respectively (Fig. S2B). Quantification of the abundance of WhiB4 per cell revealed that a major fraction of WhiB4 exists as the monomer (~ 14,000 molecules/cell), whereas the dimer (~ 4000 molecules/cell) and the trimer (~ 4000 molecules/cell) were in minor proportions under normal growth conditions (Fig. S2B and C). Pre-treatment with 0.1 mM or 0.5 mM CHP significantly increased the proportion of dimer (10,000-14,000 molecules/cell) and trimer (4000-6000 molecules/cell), with corresponding decrease of the monomer (1000-2000 molecules/cell) (Fig. S2B and C). This result is consistent with the loss of the Fe-S cluster from the holo-WhiB4 and subsequent oxidation of the exposed thiols to generate disulfide-linked oligomers of apo-WhiB4 *in vivo*. A marginal reduction in the WhiB4 levels at 0.5 mM CHP (Fig. S2B and C) is in agreement with the qRT-PCR data showing a gradual decrease in the *whiB4* transcript levels under oxidative stress (Fig. 1A). Taken together, our data indicate that oxidative stress can switch WhiB4 from a DNA binding-deficient (holo-WhiB4) to a DNA binding-proficient (oxidized apo-WhiB4) form *in vivo*.

### SI Note II

#### Redox state of WhiB4 upon overexpression

In order to understand the link between genome condensation, oxidative stress, and WhiB4, we first cloned *whiB4* into an anhydrotetracycline (Atc)-inducible plasmid containing the C-terminal FLAG epitope tag and overexpressed the protein in *MtbΔwhiB4* (*whiB4-OE*). The expression of WhiB4 in relation to Atc concentration was confirmed by western blotting (Fig. S5).

Interestingly, overexpression of WhiB4 consistently resulted in a higher proportion of dimeric and trimeric forms of oxidized apo-WhiB4 as compared to monomeric WhiB4 under normal growing conditions (Fig. S5). For example, induction of WhiB4 using 100 ng/ml of Atc generated WhiB4 monomers (~ 2000/cell), dimers (~ 14,000/cell) and trimers (~ 4000/cell) (Fig. S5), which are nearly comparable to the WhiB4 forms generated inside wt *Mtb* upon exposure to 0.1 - 0.5 mM CHP (Fig. S2).

### SI Note III

#### Functional Integrity and Physiological Relevance of WhiB4 overexpression

We tested if over-expression has any non-physiological consequences on *Mtb.* We performed qRT-PCR and measured the expression of several genes (*ahpC*, *ahpD*, *furA*, *ndh, Rv3480c, ppsC*, *blaR*) found to be overexpressed in *MtbΔwhiB4* upon CHP treatment. Indeed, the *whiB4-OE* strain retained WhiB4 activity in repressing these genes under normal growing conditions and upon CHP stress (S2 Table). As a control, the expression levels of genes not regulated by WhiB4 (*sseA, sigB, trxA, papA3,* and *ung*) were measured and found to be similar in *whiB4-OE*, *MtbΔwhiB4, and wt Mtb* (S2 Table). We also overexpressed WhiB4 without the FLAG-tag in an Atc inducible plasmid and found no significant difference between the FLAG-tagged WhiB4 and untagged WhiB4 in regulating the expression of WhiB4-specific genes (S2 Table). Previously, we had shown that *MtbΔwhiB4* has greater resistance to oxidative stress *in vitro*, inside immune-activated macrophages, and survived better in the lungs of animals, as compared to wt *Mtb* (1). In this study, we found that overexpression of WhiB4 causes growth defects specifically under oxidative stress, whereas growth was unaffected under normal growing conditions, acid stress, and heat shock (Fig. S11 A-D). Likewise, *whiB4-OE* grew poorly inside immune-activated macrophages and displayed marked attenuation in the lungs of infected mice as compared to wt *Mtb* (Fig. S11 E-F). Together, these data indicate that overexpression of FLAG tagged WhiB4 specifically reversed the phenotypic properties displayed by *MtbΔwhiB4.*

### SI Note IV

Non-lethal concentration of CHP (0.1 mM) did not induce a rapid or a significant condensation of DNA in wt *Mtb* (Fig. S9A). The mean RNS value changed from 0.25 μm ± 0.06 to 0.23 μm ± 0.06 at 6 h post-treatment (Fig. S9A). Slightly condensed nucleoids (0.17 ± 0.06 μm) were only evident at 48 h (Fig. S9A). The *MtbΔwhiB4* nucleoids underwent transitions similar to wt *Mtb*, however, the mean RNS values at 24 h and 48 h remained modestly higher than wt *Mtb* (Fig. S9A). Both wt *Mtb* and *MtbΔwhiB4* were able to maintain a similar steady-state *E_MSH_* (~-276 mV), cell length (3.0-3.5 μm), and survival at 0.1 mM CHP (Fig. S9A and S9B). Expectedly, over-expression of WhiB4 induces nucleoid condensation (mean RNS = 0.11 ± 0.02; Fig. S9A). Importantly, *whiB4-OE* displays an oxidative *E_MSH_* (~ -250 mV) under normal growing conditions, consistent with the antioxidant repressor function of WhiB4 (Fig. S9A). The oxidative shift in the *E_MSH_* of *whiB4-OE* provides a likely explanation for our earlier data demonstrating the abundance of oxidized apo-WhiB4 oligomers in *whiB4-OE* under aerobic growing conditions (as shown in figure S5). Moreover, exposure to 0.1 mM CHP further reduced the mean RNS (0.06 ± 0.01 μM), markedly increased oxidative stress (*E_MSH_*: -238 mV ± 2.85), and exerted a significant killing of the *whiB4-OE* strain (Fig. S9A and S9B).

## SI Figures

**Fig. S1:**
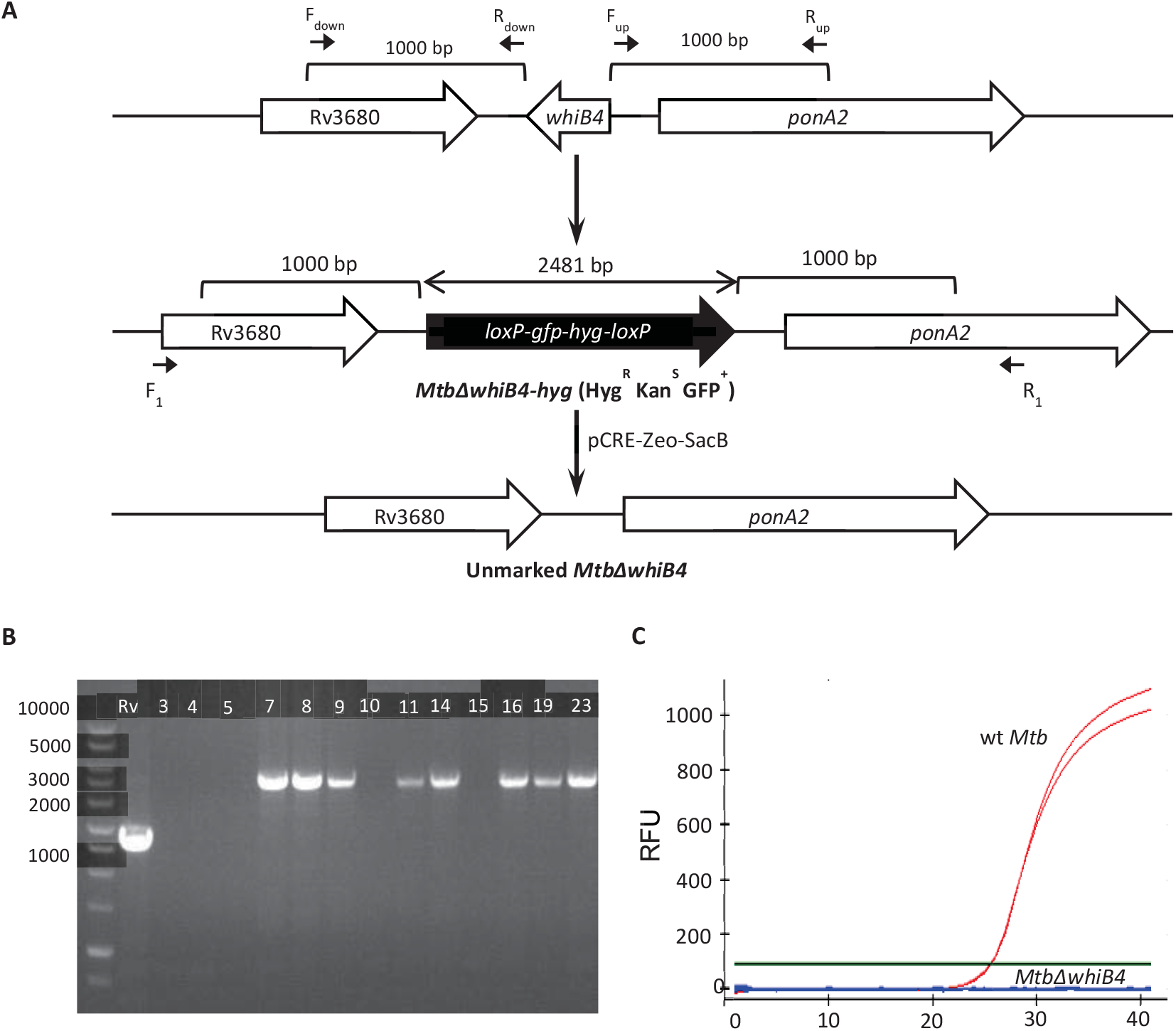
**(A)** A step-wise illustration for construction of an unmarked strain of *MtbΔwhiB4*. Upper panel denotes wt *Mtb whiB4* loci. After the allelic exchange, entire *whiB4* ORF in *Mtb* genome was replaced by right and left flanking regions of *whiB4* along with the loxP-*hyg*-*gfp*-loxP cassette. This knock out strain was unmarked by expressing Cre recombinase to remove the *hyg*-*gfp* cassette. **(B)** Genomic DNA was isolated from putative *MtbΔwhiB4* colonies (7, 8, 9, 11, 14, 16, 19 and 23) which were Kan^S^Hyg^R^GFP^+^ and replacement of *whiB4* allele with the Hyg/GFP cassette was confirmed using PCR with F1 and R1 primers (S6 Table). An increase in amplicon size from 1.2 kb to 3.3 kb due to insertion of the loxP-*hyg*-*gfp*-loxP cassette was observed in case of mutant clones, confirming the double crossover event. **(C)** RNA was isolated from logarithmically grown wt *Mtb* and the putative *MtbΔwhiB4* clones. qRT-PCR for *whiB4* was done using *whiB4* specific oligonucleotides (S6 Table) and C_t_ values were plotted to assess the expression. Amplification was detected based on fluorescence emitted by Sybergreen upon interaction with DNA and plotted as Relative Fluorescence Units (RFU).

**Fig. S2:**
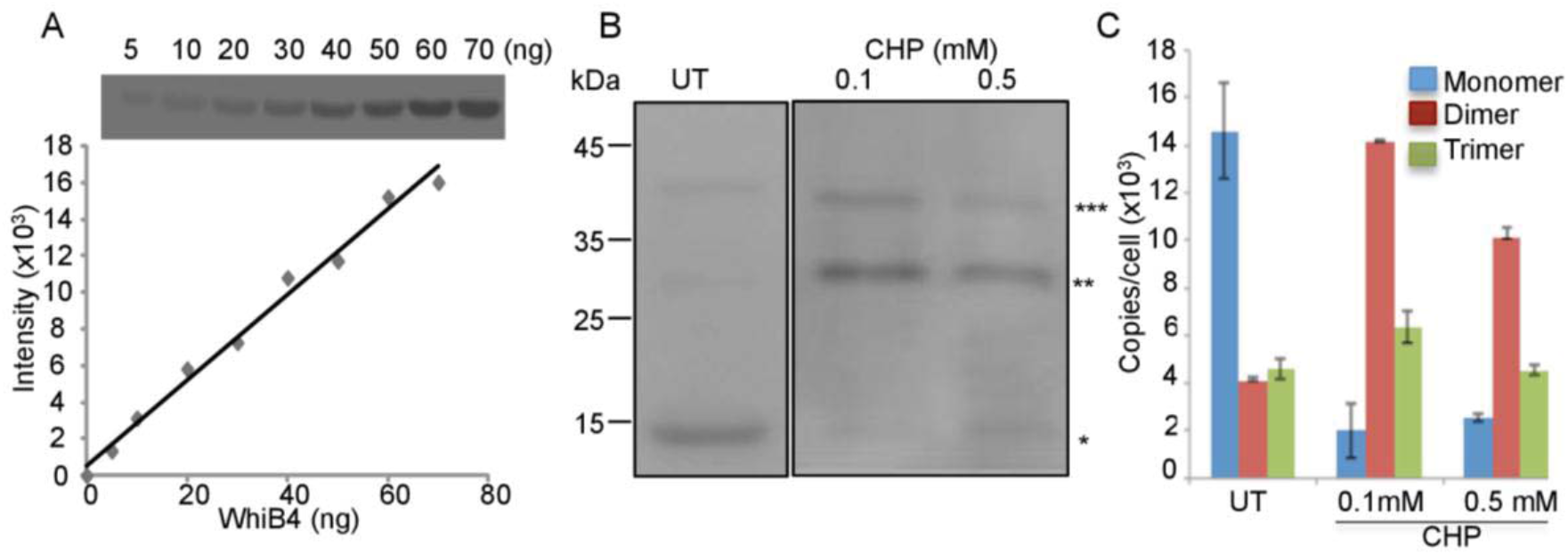
CHP stress induces the generation of oxidized apo-WhiB4 oligomers *in vivo*. (A) Indicated concentrations of purified histidine-tagged WhiB4 were subjected to immuno-blot analysis using anti-His antibody. Band intensities were quantified by Image J and plotted to generate a standard curve. (B) *MtbΔwhiB4* expressing chromosomally-integrated histidine-tagged WhiB4 from native promoter were grown till an OD_600_ nm of 0.8, harvested, and 30 μg of cell free extract was analyzed for monitoring expression and oligomerization of WhiB4 under normal growing conditions and upon exposure to 0.1 and 0.5 mM of CHP for 24 h. To minimize the possibility of O_2_-induced thiol oxidation and subsequent oligomerization of apo-WhiB4 during cell-free extract preparation, cells were pretreated with the thiol-alkylating agent NEM as described (1). Approximately 25 μg of the cell-free extract was resolved on a 12% non-reducing SDS-PAGE and immuno-blotted using anti-his. (C) The band intensities were calculated using Image J and compared with the standard curve to calculate the amount of WhiB4 expressed inside a single *Mtb* cell. These values were converted to WhiB4 copies/cell as described in Materials and Methods and plotted. Experiments were repeated at least twice in triplicate, values from both experiments were averaged and plotted.

**Figure S3:**
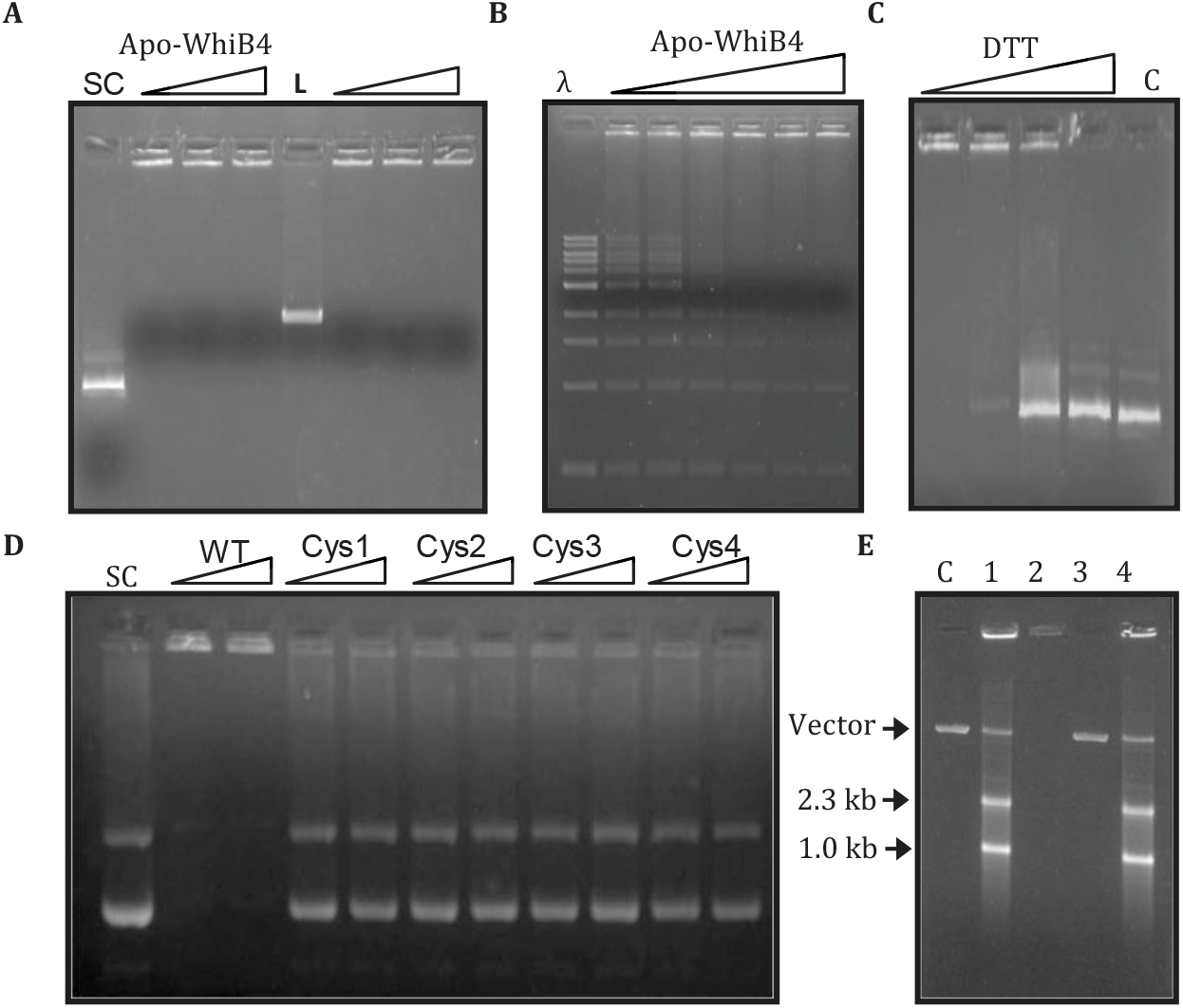
*Mtb* WhiB4 exhibits non-specific DNA binding activities. **(A)** 150 ng of either linear (L) or supercoiled (SC) plasmid DNA was incubated with oxidized apo-WhiB4 (250 ng [lanes 2, 6]; 300 ng [lanes 3, 7]; 350 ng [4, 8]). Lanes 1 and 5 represent DNA without WhiB4. **(B)** Binding of apo-WhiB4 (100, 150, 200, 250, 300 and 350 ng) with 1kb λ-DNA ladder (λ) (150 ng). **Cysteine residues regulate DNA binding of WhiB4**. **(C)** DTT-treatment abolished DNA binding of WhiB4. Oxidized apo-WhiB4 (350 ng) was treated with DTT (0, 200, 400, 800 mM) for 2 h followed by binding to the supercoiled plasmid DNA. C: represents DNA without WhiB4. **(D)** Purified wt apo-WhiB4 and Cys mutant variants were treated with thiol oxidant, diamide, and the gel-shift assay was performed as described earlier. **(E) WhiB4 represses transcription**. *In vitro* transcribed pGEM plasmid in the absence of apo-WhiB4 (lane 1), oxidized apo-WhiB4 in complex with pGEM (lane 2), *in vitro* transcription of pGEM in the presence of oxidized (lane 3) or reduced (lane 4) apo-WhiB4. The reactions in lane 3 and 4 were treated with 6 % SDS and 4 mg/ml protease K to remove WhiB4 from the mixture before separation. The samples were analyzed on 1 % agarose gel. C: pGEM plasmid DNA alone.

**Fig. S4:**
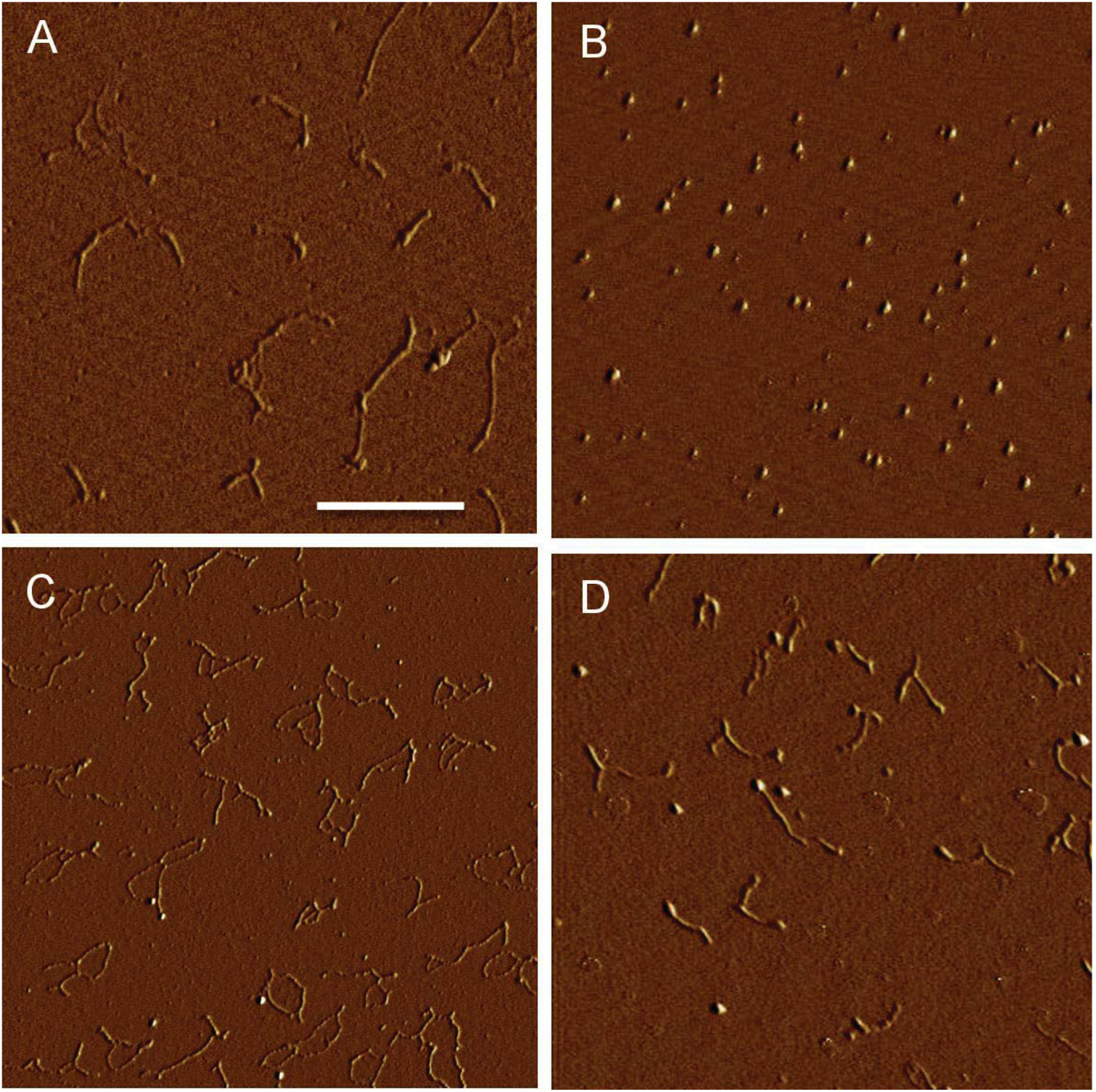
AFM image of DNA in complex with Cys3-WhiB4. **(A)** Supercoiled plasmid DNA, **(B)** Oxidized apo-WhiB4, **(C)** Oxidized apo-WhiB4 incubated with a supercoiled plasmid (at a ratio of 1:5 for DNA: Protein), **(D)** Cys3 mutant of apo-WhiB4 (Cys3-WhiB4) incubated with a supercoiled plasmid DNA at a similar ratio. The scale of images is 3 μm x 3 μm / Scale bar = 1 μm.

**Fig. S5:**
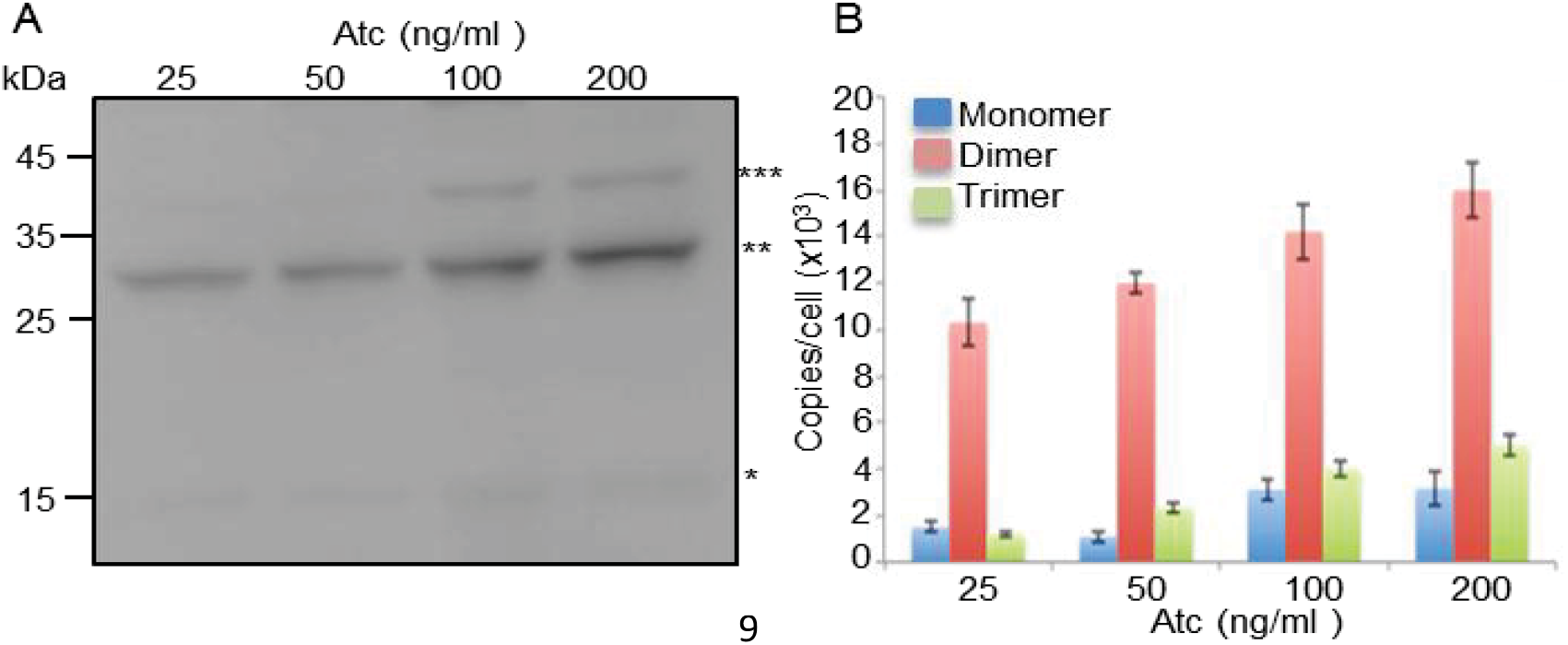
Ectopic expression of WhiB4. **(A)** *whiB4-OE* harboring FLAG-tagged WhiB4 under Atc inducible promoter was cultured till OD_600_nm of 0.4. Expression of WhiB4 was induced by addition of indicated concentration of Atc for 16h. Approximately, 30 μg of cell free extract from *whiB4-OE* strain was analyzed for WhiB4 expression by immuno-blotting using the antibody against FLAG tag. To minimize the possibility of O_2_-induced thiol oxidation and subsequent oligomerization of apo-WhiB4 during cell-free extract preparation, cells were pretreated with the thiol-alkylating agent NEM as described (1). Immuno-blot clearly revealed the appearance of monomer (*), dimers (**), and trimers (***) of WhiB4. (B) The band intensities were quantified by ImageJ software. Amount of WhiB4 expressed at a single cell level was quantified and plotted as described in figure S3.

**Fig. S6:**
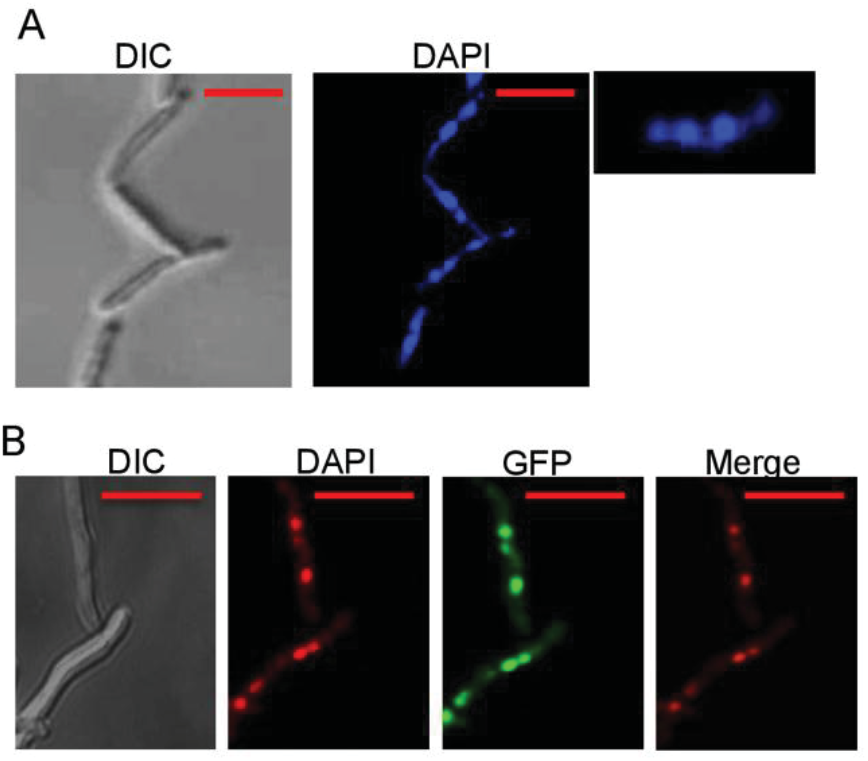
WhiB4 mediated condensation of *Mtb* genome. Confocal microscopic images (63X) of *Mtb* cells harboring (A) untagged WhiB4 under Atc inducible promoter or (B) WhiB4-GFP fusion under a strong *hsp60* promoter. In case of untagged WhiB4, expression of WhiB4 was induced by 100 ng/ml of Atc and nucleoids were stained with DAPI (pseudo colored blue). The *Inset* shows a zoomed view of a single cell showing extensive DNA condensation. In case of WhiB4-GFP fusion, nucleoids were stained with DAPI (pseudo colored red), GFP expression is indicated by green, and yellow merge indicated co-localization of WhiB4 fused GFP with DAPI stained nucleoids. *Scale bar* 2μm.

**Fig. S7:**
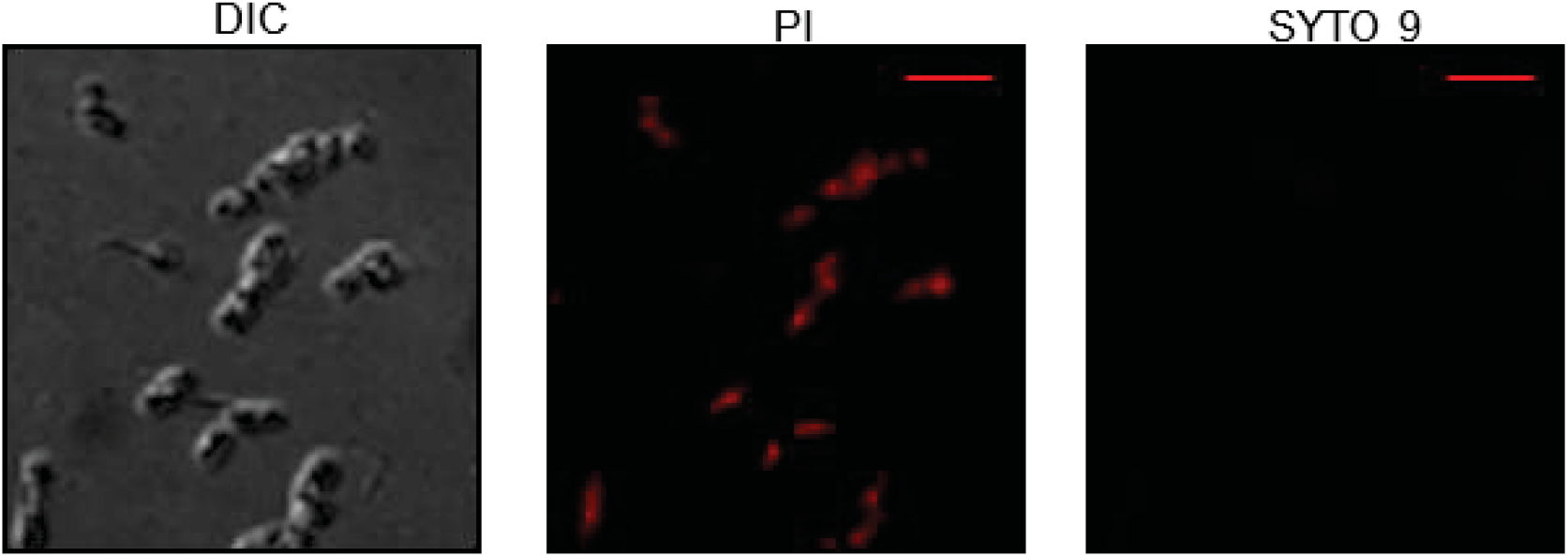
Confocal microscopic images (63X) of wt *Mtb* cells after prolonged oxidative stress. wt *Mtb* was exposed to 0.5 mM CHP treatment for 48 h and cells were stained with SYTO9 and Propidium iodide (Pi) combination. Red fluorescence emitted by Pi stained cells indicate cell death. The scale of images is 1μm.

**Fig. S8:**
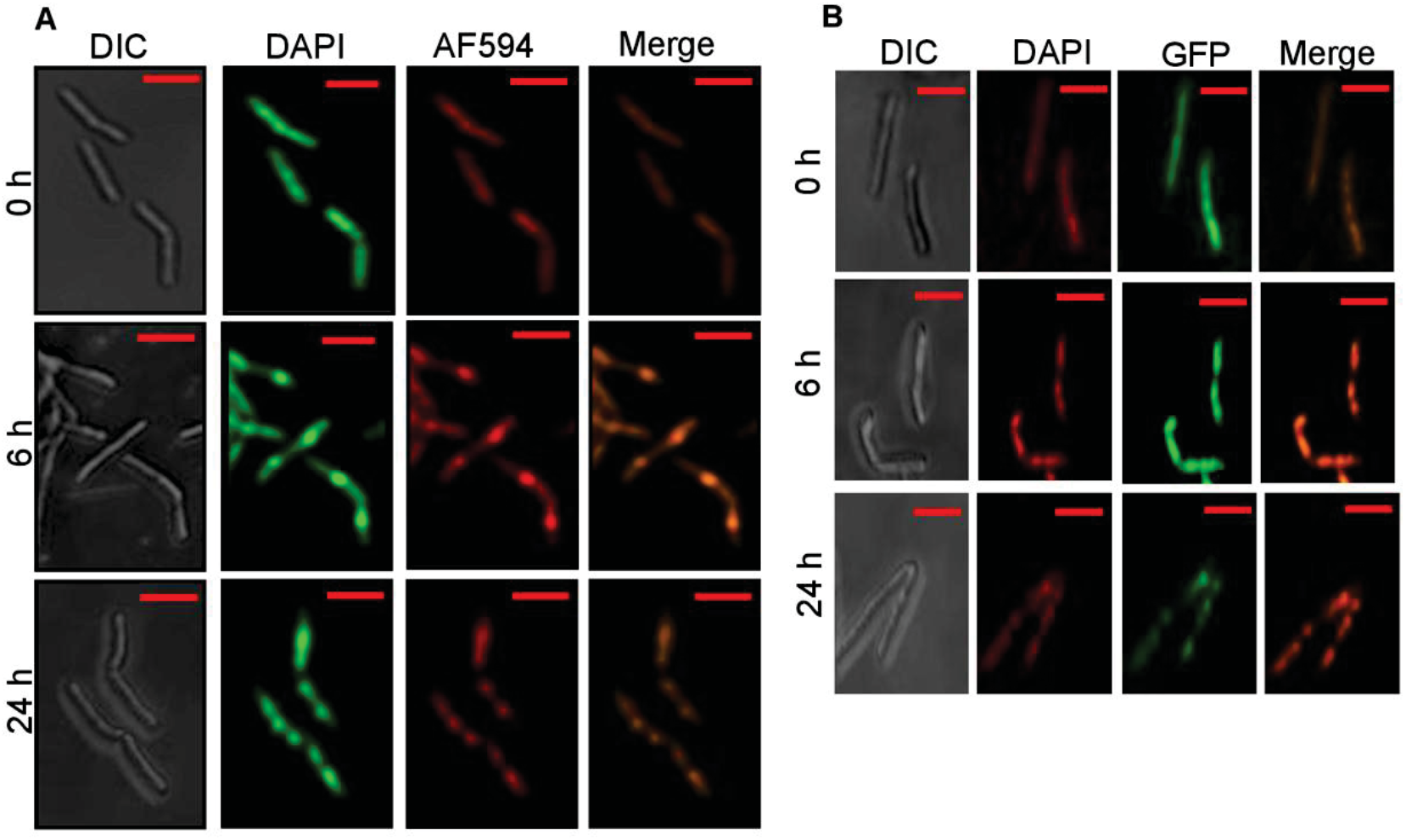
WhiB4 associates with condensed nucleoid upon oxidative stress. (A) Confocal microscopic images (63X) of *MtbΔwhiB4* cells expressing histidine-tagged WhiB4 from native promoter. The intramycobacterial *E_MSH_* of *MtbΔwhiB4* was shifted towards oxidative using 0.5 mM of CHP and WhiB4 association with nucleoids was visualized at various time points post-treatment. Nucleoids were stained with DAPI (pseudo colored green), WhiB4 was marked using anti-His primary antibody and AF594 conjugated secondary antibody (red), and yellow merge indicated co-localization of WhiB4 with DAPI stained nucleoids. **(B)** Confocal microscopic images (63X) of *Mtb* cells expressing GFP fused WhiB4 in presence of 0.5 mM CHP for the given time points. Nucleoids were stained with DAPI (pseudo colored red), GFP expression indicated by green, and yellow merge indicated co-localization of WhiB4 fused GFP with DAPI stained nucleoids. The scale bar is 2 μm.

**Fig. S9:**
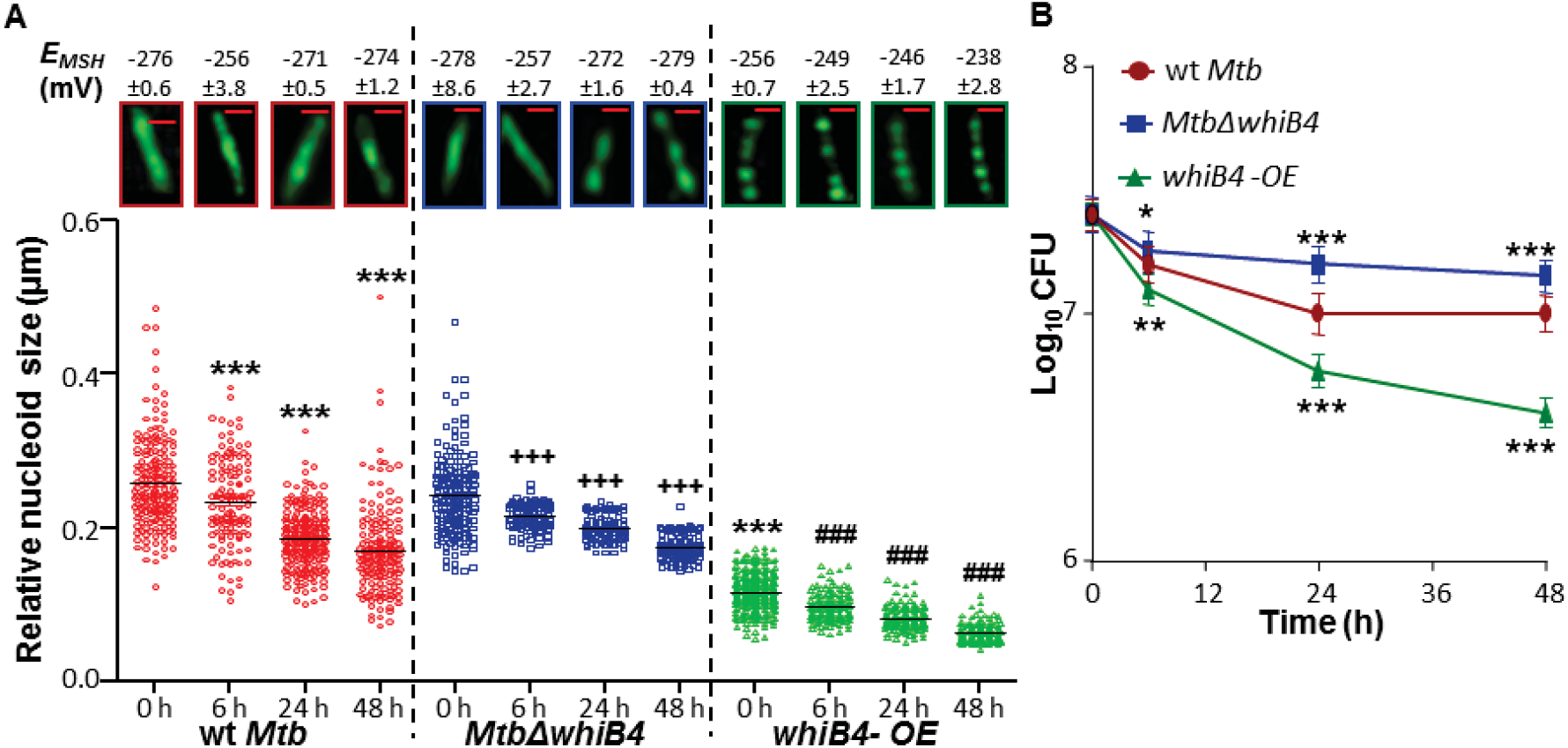
Oxidative stress leads to DNA condensation, skewed redox homeostasis and bacterial killing in a WhiB4-dependent manner. (A) wt *MtbΔwhiB4* and *whiB4-OE* were exposed to 0.1 mM CHP for 6, 24 and 48 h. Nucleoids were stained with DAPI (pseudo colored green) and visualized by confocal microscopy (63X). The scale of images is 1μm. For inducing WhiB4, 100 ng/ml of Atc was added to cultures of *whiB4-OE*. As a control, the same amount of Atc was added to other strains. The relative RNS of ~ 100-150 independent cells was measured and represented as scatter plots. Each dot represents one nucleoid. The line depicts mean of the population at each time point. *P values*: * = as compared to *wt Mtb,* + = as compared to *MtbΔwhiB4,* # = as compared to *whiB4-OE* at 0 h (\*\*\**P* or ^+++^*P* or ^###^*P* ≤ 0.001). Intramycobacterial *E_MSH_* of various *Mtb* strains at each time point was measured using flowcytometer and shown along with a corresponding image of a representative DAPI stained cell visualized by confocal microscopy under similar conditions. **(B)** Plots showing the survival of *Mtb* strains in response to CHP stress as assessed by enumerating CFUs. Data shown is the average of experiments performed in triplicates. Error bars represent SD from the mean.

**Fig. S10:**
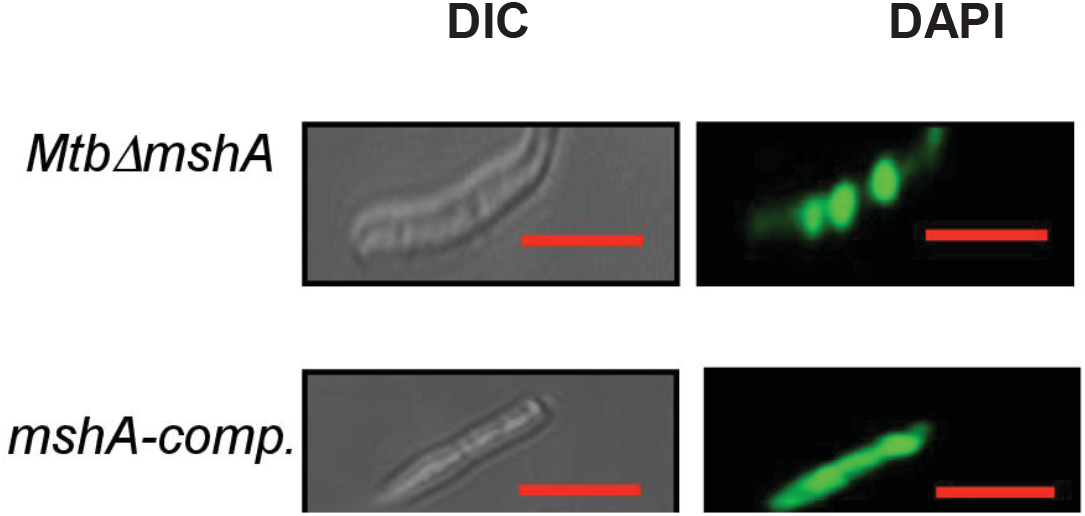
Condensation of *Mtb* genome upon Mycothiol depletion. **(A)** A representative image of nucleoids of *MtbΔwhiB4* and *mshA-comp* strains. Nucleoids were stained with DAPI (pseudo colored green) and visualized under the confocal microscope (63X). The mean RNS values of ~ 100-150 cells of *MtbΔwhiB4* and *mshA-comp* strains were 0.11 ± 0.01 μm and 0.25 ± 0.15 μm, respectively. The scale of images is 3 μm.

**Fig. S11:**
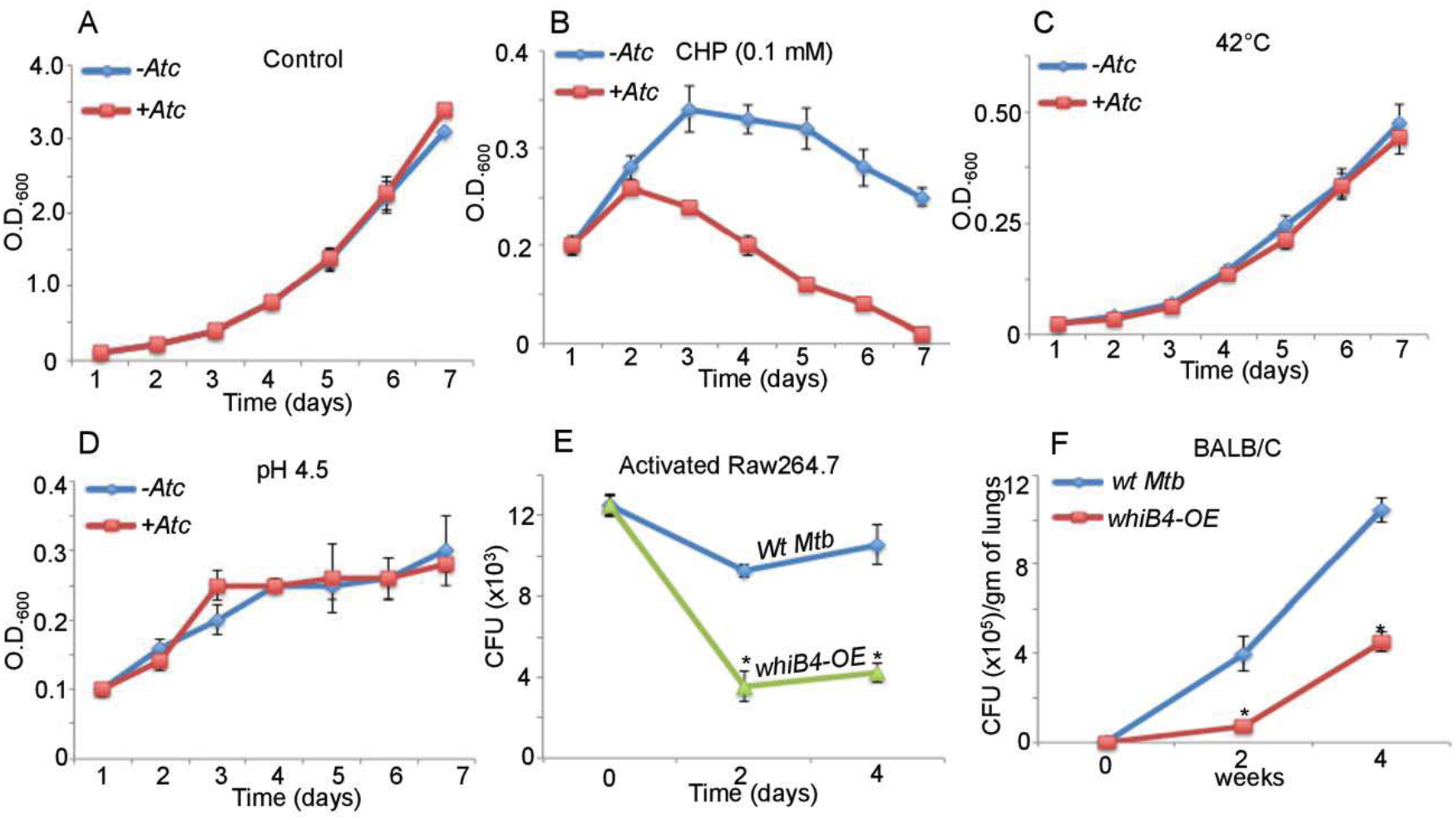
WhiB4 overexpression specifically reverses phenotypes displayed by *MtbΔwhiB4*. We have earlier shown that *MtbΔwhiB4* survived better upon oxidative stress *in vitro*, inside immune-activated macrophages, and displayed hypervirulence in animals (1). To investigate the functional relevance of WhiB4 overexpression, we determined the phenotype of WhiB4-OE under defined conditions specific to the WhiB4 function. The *whiB4-OE* strain was either subjected to 200 ng/ml of Atc induction or left uninduced. Cells were monitored for growth at different time intervals by measuring absorbance at 600 nm under **(A)** aerobic conditions, **(B)** 0.1 mM CHP, (C) 42ºC, and **(D)** pH 4.5. Data shown is the average of two experiments performed in triplicates. Error bars represent SD from the mean. **(E)** IFN-Ɣ-LPS activated RAW 264.7 macrophages were infected with wt *Mtb*, and *whiB4-OE* strains at an MOI of 10 and survival was monitored by enumerating CFU. Infected macrophages were maintained in 200 ng/ml of Atc for sustained induction of WhiB4. The data is shown as Mean (± SD) from two independent experiments having triplicate wells each. *P < 0.05. **(F)** BALB/c mice were infected by aerosol and survival of various strains was monitored at 2 and 4 week’s time post infection in lungs. Error bars represent SD from the mean. To ensure the over-expression of WhiB4, doxycycline dissolved in water was supplied in the drinking water for mice. The concentration of doxycycline was maintained at 1 mg/ml in 5% sucrose solution.

**Fig. S12:**
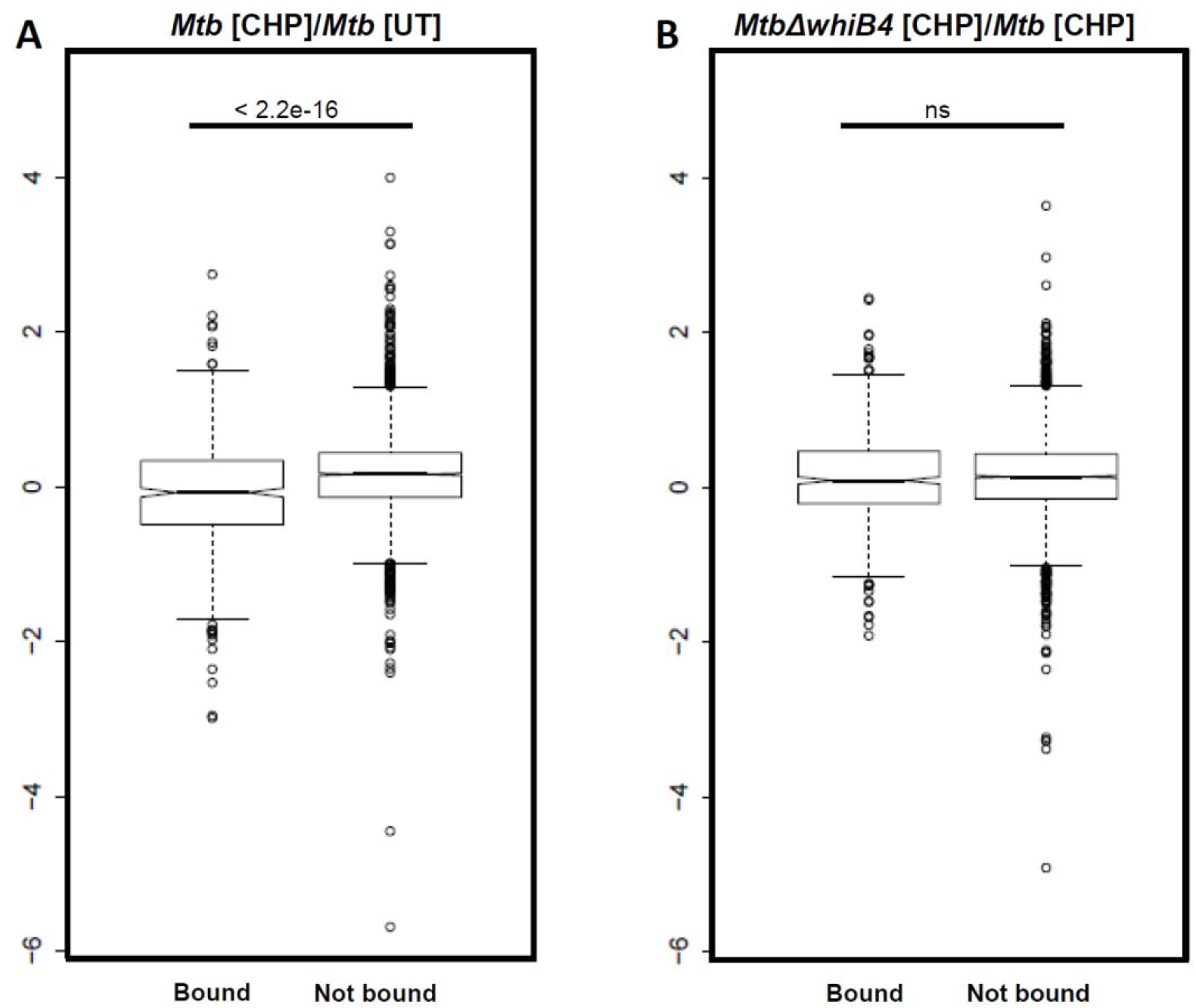
Comparison of Chip-Seq binding data for WhiB4 with gene expression changes for (A) wt *Mtb* CHP vs. wt *Mtb* (B) *MtbΔwhiB4* CHP vs wt *Mtb* CHP. Gene expression log-2 fold change of bound and not bound genes.

**Table S1:** Differentially regulated genes in microarray in wt *Mtb* and *MtbΔwhiB4* in response to CHP [(log values) 2-fold up- and down-regulation, p≤0.05]. (Enclosed as a separate excel spread sheet).

**Table S2:** qRT-PCR analysis of CHP responsive genes.

| Gene | wt *Mtb* (T) vs<br>wt *Mtb*(UT) | *MtbΔwhiB4*<br>(T) vs wt<br>*Mtb*(T) | *whiB4-OE*<br>(UT) vs<br>*MtbΔwhiB4*<br>(UT) | *whiB4-OE*<br>(T) vs<br>*MtbΔwhiB4*<br>(T) | *whiB4-OE*<br>(-FLAG[T])<br>vs<br>*MtbΔwhiB4*<br>(T) |
| --- | --- | --- | --- | --- | --- |
| *ahpC* | 2.65±0.47<br>(4.34) | 2.26±0.17<br>(2.30) | -3.85±0.49 | -10.67±1.52 | -7.56±0.87 |
| *ahpD* | 3.31±0.61<br>(5.11) | 2.04±0.03<br>(2.09) | -6.66±0.36 | -9.24±0.29 | -11.24±2.6 |
| *cydA* | 1.8±0.60<br>(1.85) | 3.23±0.31<br>(2.59+2.43) | -3.90±0.17 | -4.65±0.17 | -2.97±0.13 |
| *furA* | 11.31±1.73<br>(20.14) | 2.55±0.35<br>(1.95) | -2.79±0.40 | -6.49±0.49 | -3.04±1.43 |
| *ppsC* | 1.65±0.81<br>(1.02) | 4.84±0.77<br>(5.24) | -2.43±0.16 | -2.65±0.05 | -4.57±1.87 |
| *ppsD* | -1.34±0.10<br>(1.20) | 3.63±0.55<br>(2.65+2.69) | -3.54±0.77 | -6.49±0.49 | -5.13±1.78 |
| *blaR* | 2.19±0.22<br>(1.07) | 4.65±1.47<br>(2.73) | -4.95±1.17 | -2.65±0.47 | -3.23±1.1 |
| *Ndh* | 5.36±0.81<br>(5.04) | 6.19±0.98<br>(4.24) | -2.05±0.02 | -5.15±1.97 | -4.61±1.3 |
| *Rv3480c* | 1.18±0.47<br>(-1.58) | 3.00±0.02<br>(1.71) | -2.95±1.03 | -3.65±0.72 | -1.97±0.31 |
| *\*papA3* | -1.1±0.111<br>(-1.03) | 1.06±0.07<br>(-1.71) | -1.13±0.31 | -1.1±0.08 | 1.3±1.08 |
| *\*trxA* | 1.356±0.034<br>(1.48) | -1.25±0.03<br>(-1.17) | 1.15±0.03 | -1.04±0.47 | -1.34±0.56 |
| *\*sigB* | 5.24±0.03<br>(4.19) | 1.07±0.07<br>(1.09) | -1.31±0.16 | -1.32±0.28 | 1.11±0.12 |
| *\*sseA* | 1.103±0.019<br>(1.29) | -0.89±0.017<br>(-1.11) | 1.10±0.23 | -0.69±0.13 | -1.69±1.34 |
| *\*ung* | -2.30±0.42<br>(2.15) | 1.51±0.13<br>(-1.22) | 1.12±0.09 | -1.65±0.17 | -0.5±0.09 |
| *\*mutY* | -2.26±0.08<br>(-1.90) | 1.54±0.44<br>(-1.23) | -1.26±0.08 | -1.13±0.14 | 1.32±0.14 |

Values given in red are the absolute fold changes in microarray data with the (-) sign depicting down-regulation. “\*” Denotes genes not regulated by WhiB4 under CHP stress. CHP exposed samples were denoted as “T”. The *whiB4-OE (-FLAG)* strain expresses an untagged version of WhiB4 from an Atc inducible promoter.

**Table S3A: Raw Chip-seq data showing enrichment of WhiB4 on regions of *Mtb* chromosome.** (Enclosed as a separate excel spread sheet).

**Table S3B: Comparison of Chip-seq binding data for WhiB4 with expression changes obtained for wt *Mtb* CHP vs. untreated wt *Mtb*.** (Enclosed as a separate excel spread sheet).

**Table S3C: Comparison of Chip-seq binding data for WhiB4 with expression changes obtained for** MtbΔwhiB4 **CHP vs Wt Mtb CHP.** (Enclosed as a separate excel spread sheet).

**Table S4:**
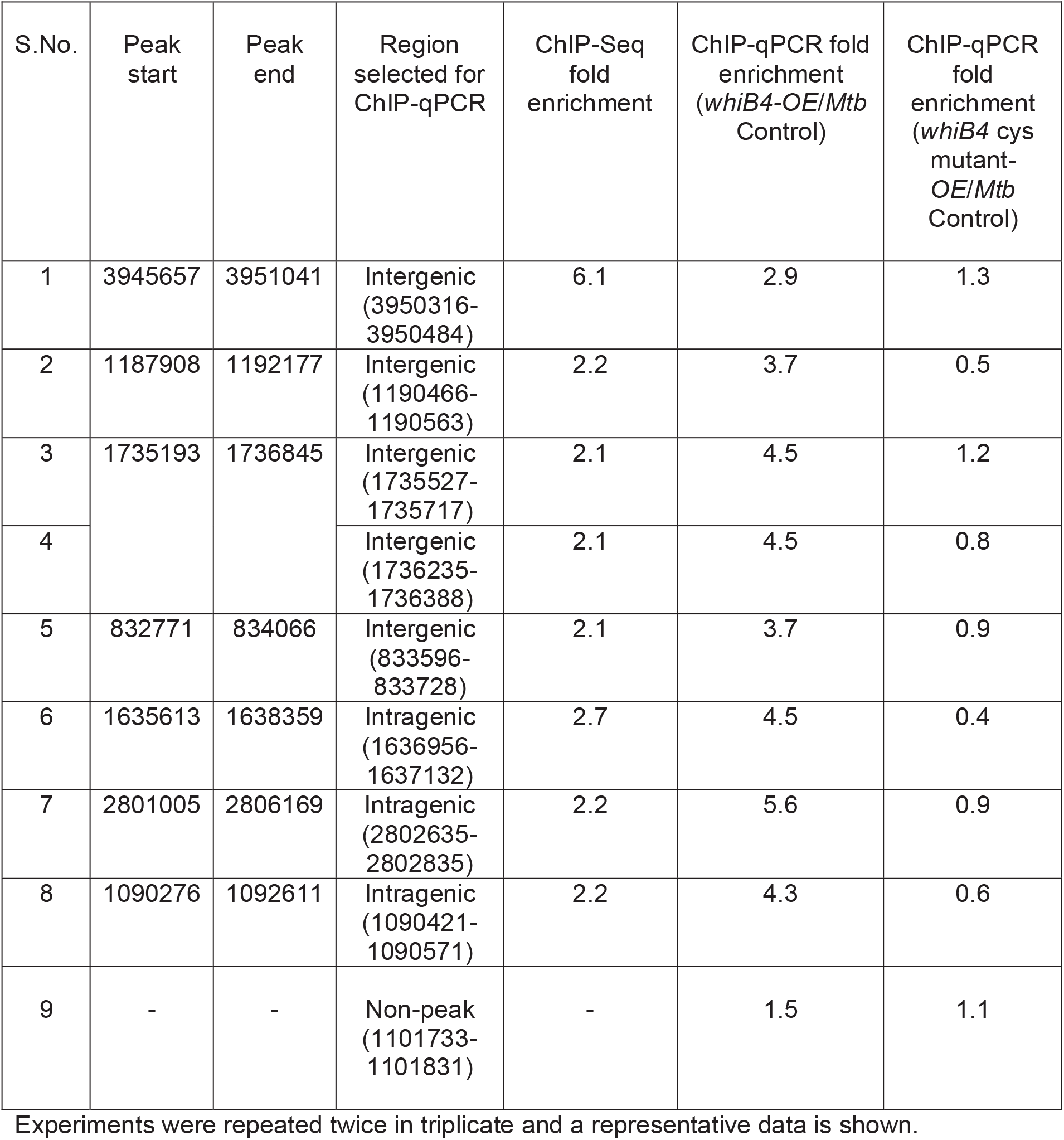
ChiP-qPCR to validate ChIP-seq.

**Table S5:** *Mtb* strains used in this study.

| Strain | Description |
| --- | --- |
| *wt Mtb H37Rv* | *Mycobacterium tuberculosis* wild type parental strain |
| *MtbΔwhiB4* | *whiB4* knockout strain of *Mtb* |
| *whiB4-Comp* | *whiB4* complemented strain of *Mtb* |
| FLAG tagged *whiB4-OE* | FLAG tagged *whiB4* overexpressing strain of *Mtb* |
| *whiB4-OE* | Untagged *whiB4* overexpressing strain of *Mtb* |
| *whiB4-cys3-OE* | *whiB4* cys3 mutant overexpressing strain of *Mtb* |
| *MtbΔmshA* | *mshA* knockout strain of *Mtb* |
| *mshA-comp* | *mshA* complement strain |
| *HU-OE* | *hu* overexpressing strain of *Msm* |

**Table S6:**
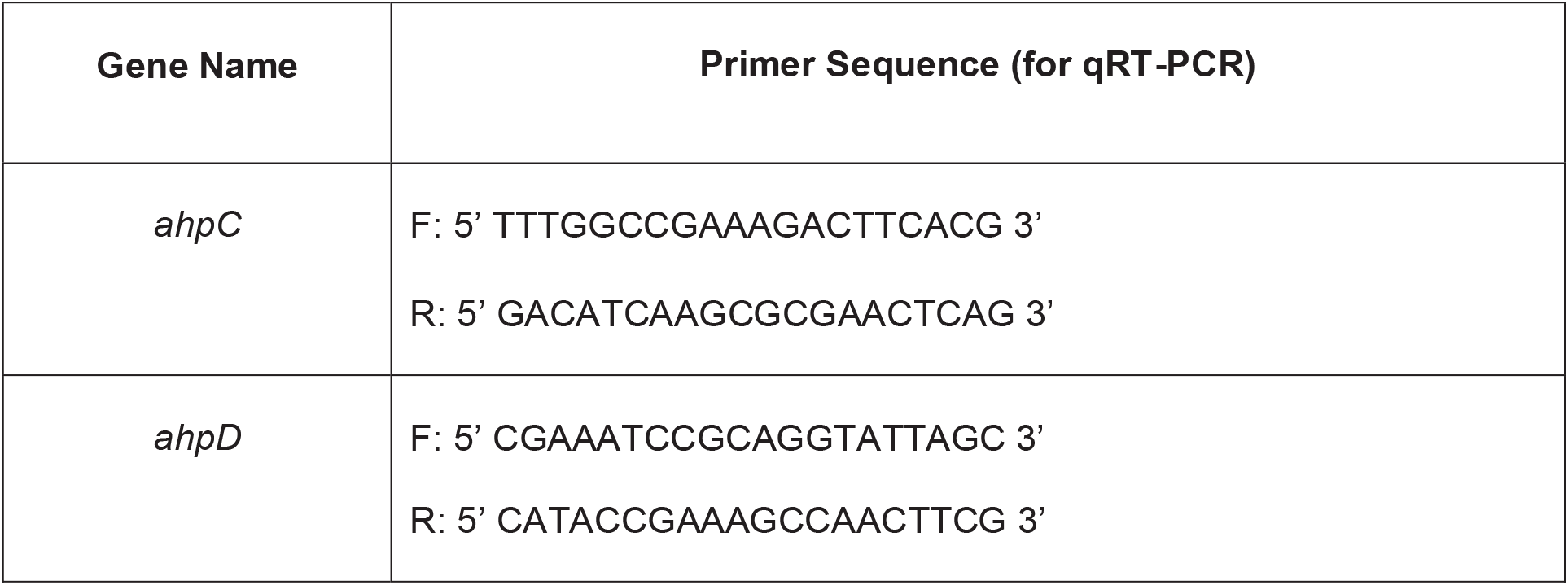
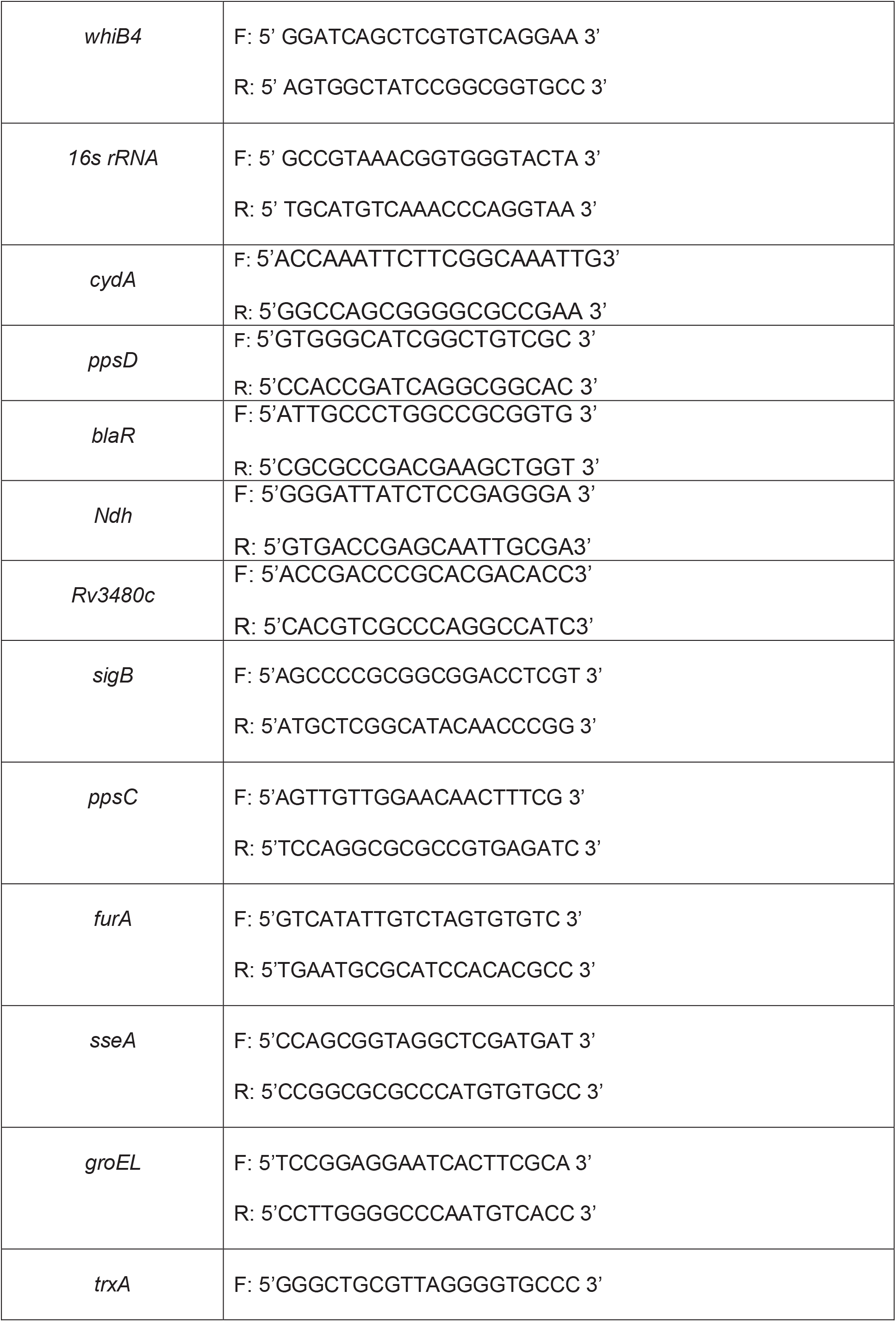
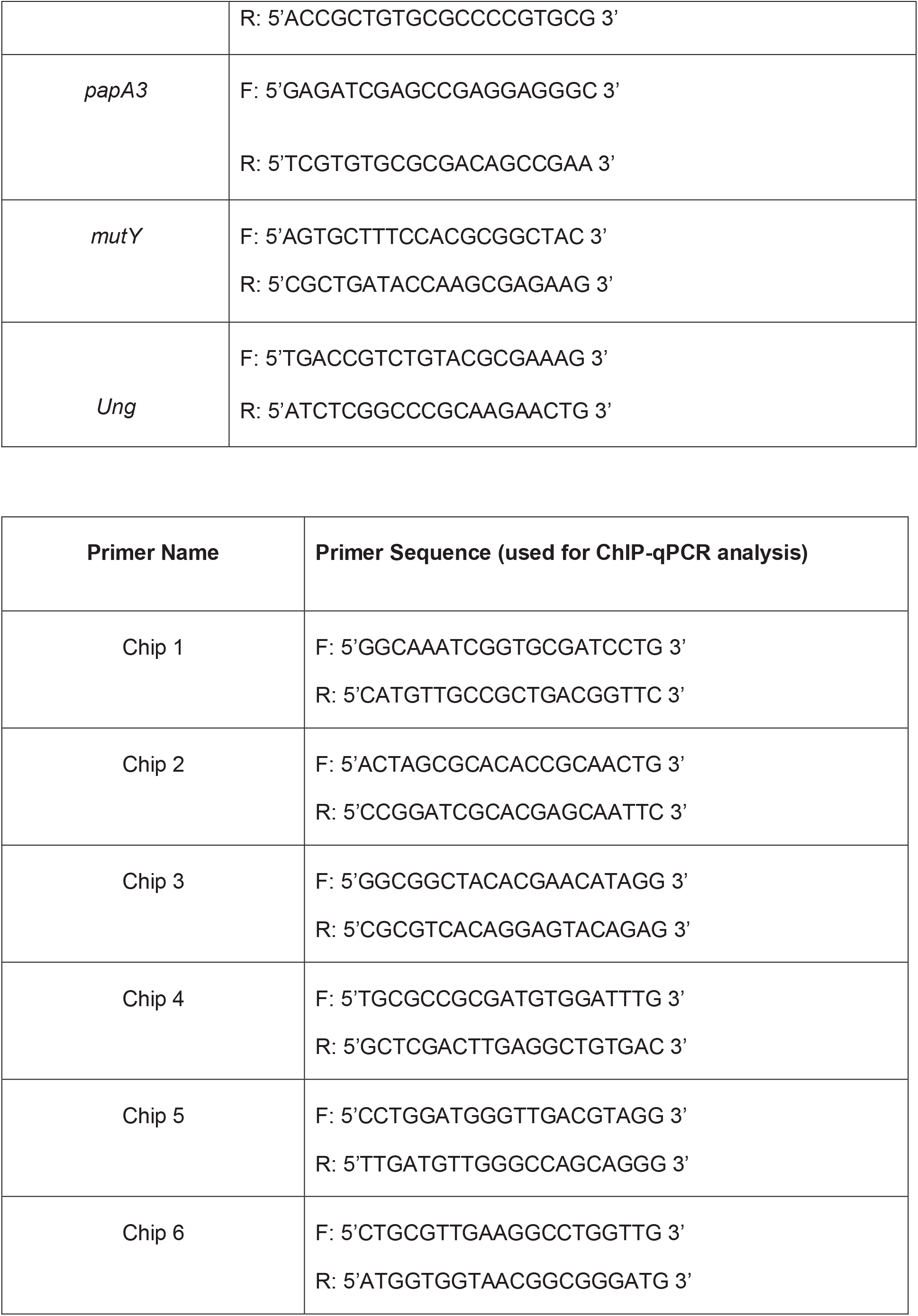
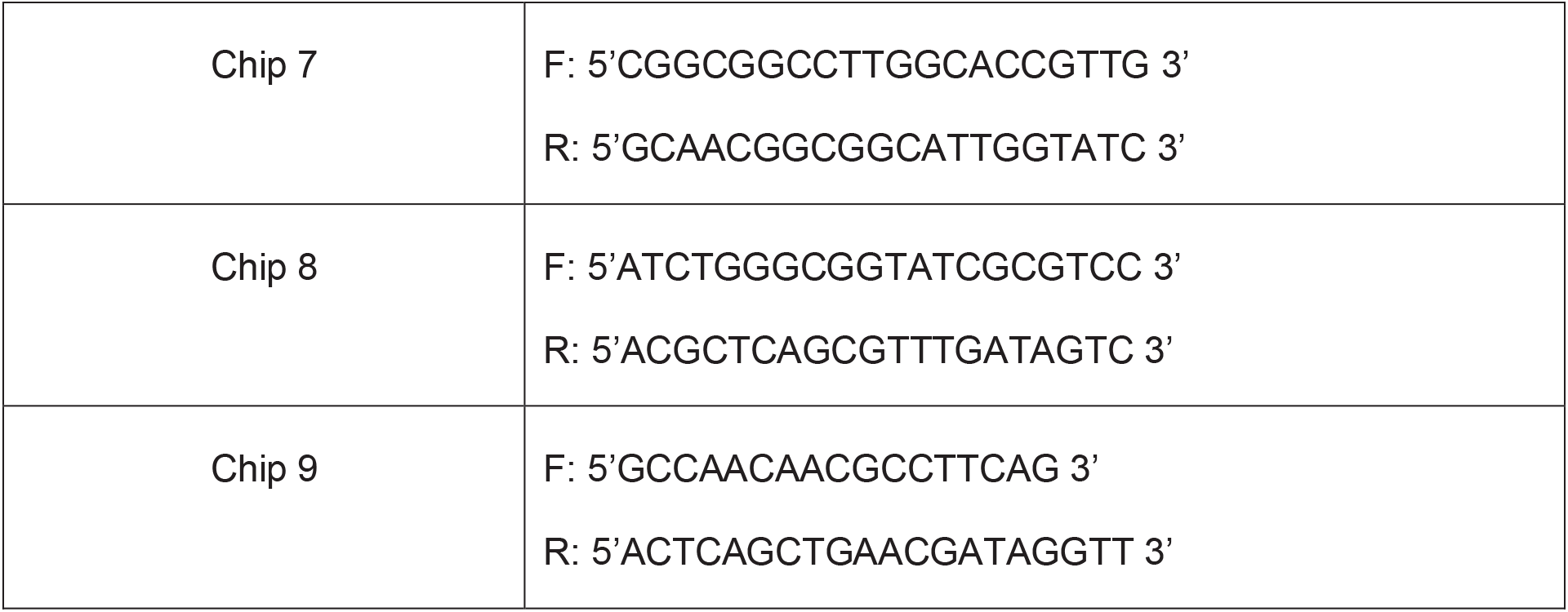
Sequences of primers used in this study.

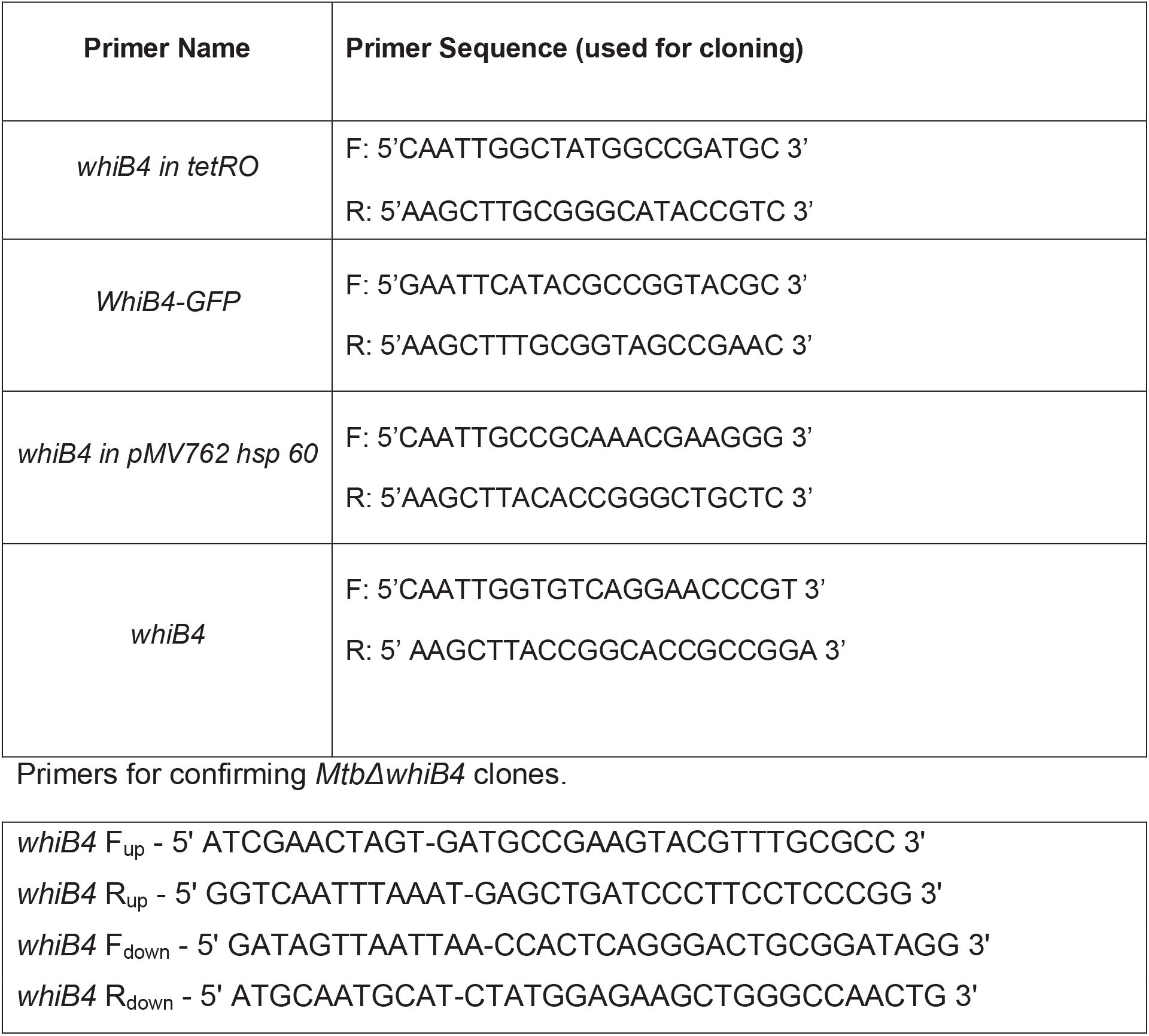
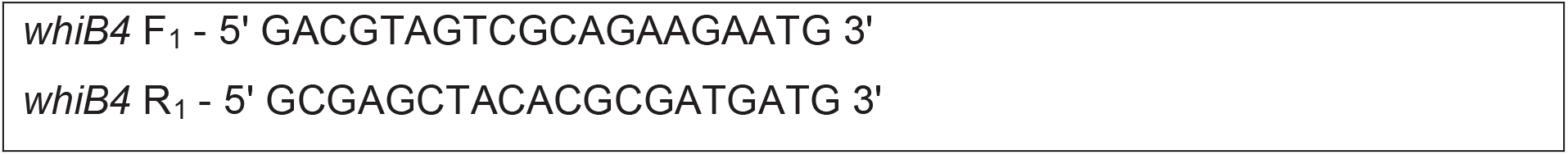
Primers for cloning whiB4 in tetRO and cloning of WhiB4-GFP fusion.

## SI Materials and Methods

### Microarray experiments

The wt *Mtb* and *MtbΔwhiB4* strains were grown in 7H9 medium supplemented with 1x ADS (Albumin Dextrose Saline enrichment) to an O.D._600_ of 0.4 and exposed to 0.25 mM CHP for 2 h at 37^o^C. The experiment was carried out with two biological replicates of both the strains. At 2 h post-treatment with CHP, total RNA was purified as described (1). The total RNA was processed and hybridized to the *Mtb* Whole Genome Gene Expression Profiling microarray-G2509F (AMADID: G2509F_034585, Agilent Technologies PLC). DNA microarrays were provided by the University of Delhi South Campus MicroArray Centre (UDSCMAC). RNA amplification, cDNA labeling, microarray hybridization, scanning, and data analysis were performed at the UDSCMAC as described previously (2). Slides were scanned on a microarray scanner (Agilent Technologies) and analyzed using the GeneSpring software. Results were analysed in MeV and data was considered significant at p ≤ 0.05. The normalized data has been submitted to NCBI’s Gene Expression Omnibus (GEO, <u>http://www.ncbi.nlm.nih.gov/geo/</u>); accession number GSE73877.

### qRT-PCR

Total RNA was isolated from logarithmically grown cells of wt *Mtb*, *MtbΔwhiB4 and whiB4-OE* strains and qRT-PCR was performed using gene specific primers (Table S6) as described (1).

### Electrophoretic Mobility Shift Assays (EMSA)

Histidine-tagged version of WhiB4 and its various cysteine mutants were purified by as described elsewhere (1). The oxidized apo-form of WhiB4 was prepared as described previously (1). To assess the non-specific DNA binding ability of WhiB4, oxidized apo-WhiB4 was incubated with various DNA molecules in a buffer containing 89 mM Tris, 89 mM boric acid and 1 mM EDTA (pH 8.4) for 30 min. The protein-DNA complexes were resolved on a 1% agarose gel in 1x TBE buffer, stained with ethidium bromide and visualized under UV light.

### *In vitro* transcription assays

Approximately 350 ng of oxidized apo-WhiB4 was incubated with 150 ng of a pGEM plasmid in a final volume of 20 μl, followed by *in vitro* transcription using the Riboprobe *in vitro* Transcription Kit according to the manufacturer’s instructions (Promega). Samples were treated with 6% SDS (Amresco Inc.) and 4 mg/ml proteinase K (Amresco Inc.) for 30 min at 37^o^C and the reaction products were analyzed on a 1% agarose gel.

### *In vivo* thiol-trapping analysis

Histidine-tagged WhiB4 was expressed under its native promoter in *MtbΔwhiB4*. At an O.D._600_ of 0.4, *MtbΔwhiB4* expressing histidine-tagged WhiB4 was treated with 0.1 and 0.5 mM CHP for 24 h. Cultures were treated with 10 mM NEM for 5 min to block the thiol-state of WhiB4. Bacterial pellets were resuspended in lysis buffer [300 mM NaCl, 20 mM Na-Phosphate, 10% Glycerol and 1X protease inhibitor (Amresco Inc.), pH 7.5] and lysed using bead beater (MP Biomedicals). Approximately 30 μg of total cell lysate was resolved by 12% non-reducing SDS-PAGE. Proteins were transferred on to 0.2 μm PVDF membrane and used for Western blot. Western blot analysis was achieved using 1:20,000 dilution of anti-His antibody (GE Life Sciences) for 12 h. The blotted membrane was developed with a 1:20,000 dilution of peroxidase-conjugated anti-mouse IgG (Cell Signaling) and an enhanced chemiluminescence substrate (GE Amersham).

### Immuno-blot analysis

The expression of WhiB4 in the *whiB4-OE* and *whiB4-cys3-OE* strains was induced using various concentrations of Atc for 16 h. For the immuno-blot assay, bacterial cells were processed as described earlier. WhiB4 was detected using anti-FLAG primary antibody. The blot was developed with enhanced chemiluminescence (ECL, Bio-Rad).

### Confocal microscopy

Various *Mtb* strains were grown to exponential phase (O.D._600_ of 0.4) in 7H9 medium and WhiB4 expression was induced as described (1). Cells were fixed with paraformaldehyde (PFA) and washed with 1X PBS and stained with 4’,6-diamino-2-phenylindole (DAPI, 1 μg/ml, Invitrogen). The bacterial cells were visualized for DAPI fluorescence (excitation at 350 nm and emission at 470 nm) in a Leica TCS Sp5 confocal microscope under a 63X oil immersion objective. Quantification of nucleoid size (larger axis) of 100-150 independent cells was carried out using public domain program OBJECT-IMAGE J. Using these images, relative nucleoid size (RNS) was measured by determining the ratio between length of the nucleoid(s) and the length of the cell by relying on the end points of their larger axes. We only considered cells with bilobed or irregular shaped nucleoid for measurements. For this measurement, we assumed that the width of the nucleoid and that of *Mtb* cells are similar with no significant differences. We observed be minor variations in cell size and shape among different *Mtb* strains. Since the images are two-dimensional and *Mtb* cells are small, the value of RNS are merely approximations indicating the trend of compactions or expansion of the nucleoid under conditions studied. For each condition 100-150 independent cells were analyzed using the public domain program OBJECT-IMAGE J, and the data were compiled and analyzed to obtain a mean RNS value with error bars.

### Immunofluorescence microscopy

The *whiB4-OE* strain was fixed with 4% PFA, washed with 1X PBS and permeabilized using 0.1% of Triton-X 100. Cells were blocked with 2% bovine serum albumin (BSA) and incubated with anti-FLAG or anti-His primary antibody. The cells were incubated with anti-mouse IgG Alexa Fluor® 594-conjugated secondary antibody (Invitrogen) followed by staining with DAPI as described earlier. A Leica TCS Sp5 confocal microscope under a 63X oil immersion objective was used for imaging.

### Live dead cell staining

CHP treated cells were washed with 1X PBS and stained using the Live/Dead BacLight Bacterial Viability Kit (Invitrogen). The cells were fixed with 4% PFA and imaged for SYTO9 dye (excitation at 480nm and emission at 500nm) and propidium iodide (excitation at 490nm and emission at 635nm) in a Leica TCS Sp5 confocal microscope.

### ChIP-Seq

DNA–protein interactions were characterized by cross-linking 100 ml of culture of wt *Mtb* or *whiB4-OE* strains (O.D._600_ of 0.4-0.5) with 1% formaldehyde while agitating cultures at 37^o^C for 30 min. Crosslinking was quenched by the addition of glycine to a final concentration of 125 mM. Cells were pelleted, washed in 1X PBS+1X protease inhibitor cocktail (Sigma), and resuspended in IP Buffer (20 mM K-HEPES pH 7.9, 50 mM KCl, 0.5 mM dithiothreitol and 10% glycerol) + the 1 X protease inhibitor cocktail. Cell debris was removed by centrifugation after lysis using Bioruptor and the supernatant was used in the IP experiment. The samples were mixed with buffer IPP150 (10 mM Tris-HCl—pH 8.0, 150 mM NaCl and 0.1% NP40) and immunoprecipitation of FLAG-tagged proteins was initiated by incubating samples overnight rotating at 4 °C with Anti-FLAG® M2 Magnetic Beads (Sigma: M8823). Beads were washed twice with IP buffer and once with Tris-EDTA buffer pH 7.5. Elution was performed in 50 mM Tris–HCl pH 7.5, 10 mM EDTA, 1% SDS for 40 min at 65°C. Samples were finally treated with RNAse A for 1 h at 37°C, and cross-links were reversed by incubation for 2 h at 50°C and for 8 h at 65°C in elution buffer with Protease K. DNA was purified by phenol-chloroform extraction and quantified. The concentration of the immunoprecipitated DNA was measured using the Qubit HS DNA kit. The resulting ChIP-DNA was subjected to qRT PCR analysis to determine the enrichment in the immunoprecipitated sample (*whiB4-OE*) over the control sample (wt *Mtb*) after normalization with input. Truseq ChIP kit was used for the library preparation. DNA was end-repaired and adapters were ligated. After ligation, products were purified and amplified. Samples were sequenced on the HiSeq 2500 Sequencing System.

### ChIP-Sequencing analysis

The single end reads obtained were aligned to the reference genome H37Rv (ASM19595v2) using the bwa-0.7.12 version. The tab delimited *.sam* files as output were then converted to .bam format using samtools-1.2. We used bedtools2 to convert .bam into .bed files, and both the resultant ChIP and input files in replicates were used in MACS peak calling algorithm (version2). The p-value cutoff for the peaks was set to be 0.01. Peaks obtained for each condition were visualized in the UCSC microbial genome browser. The analysis was also performed using the Z-score approach (3, 4). Read counts for each base position were calculated using bedtools2 coverage command. This approach assumes normal distribution of the read counts and firstly normalizes the ChIP samples and the corresponding input controls individually with the mode of the distribution. Mode was calculated using the Shorth function in Genefilter package in R. The cut-off Z-score value was then used as the threshold to filter the base positions. For further analysis, only positions that were present in both the biological replicates were considered for further analysis. The adjacent positions as well as distance less than 50 base pairs were merged. We compared the peaks obtained from both the methods and found ~90% similarity. The %GC was calculated using bedtools2 nuc command, with the default parameters. All the related statistical analyses were performed in R. Peak coordinates were then intersected with the annotated H37Rv file. Peaks that were found within 300 bp upstream and 50 bp downstream of the start sites of the genes were inferred as bound. In parallel, genes were categorized as up-regulated (>= 1.5 log_2_ fold change) and down-regulated (<= -1.5 log_2_ fold change) from the microarray datasets. These bound genes were then overlapped with the up- and down-regulated genes to understand the effect of WhiB4 binding on the regulatory regions and the downstream effects on gene expression.

### Macrophage and mouse infections

IFN-Ɣ+LPS activated RAW 264.7 macrophages were infected with wt *Mtb* and *whiB4-OE* strains at a multiplicity of infection 10 (MOI 10) of 2 for 4 h, followed by treatment with amikacin to remove extracellular bacteria. After infection, cells were washed thoroughly with warm DMEM medium and resuspended in the same containing 10% FBS. Samples were collected at the indicated time points, lysed using 0.06% SDS-7H9 medium and plated on OADC-7H11 agar medium for CFU determination.

BALB/c mice were infected via aerosol (Wisconsin-Madison) with the wt *Mtb* and *whiB4-OE* strain. To ensure the over-expression of WhiB4, doxycycline dissolved in the drinking water. The concentration of doxycycline was maintained at 1 mg/ml in 5% sucrose solution. At selected time point, three mice were killed and their lungs were removed and processed for analysis of bacillary load. The number of CFUs was determined by plating appropriate serial dilutions on 7H11 plates. Colonies were counted 3-4 weeks of incubation at 37°C.

### Statistical analysis

The overlap was assessed using the GeneOverlap package within R package version 1.6.0 (<u>http://shenlab-sinai.github.io/shenlab-sinai/</u>) and statistical significance was calculated using the Fisher’s exact test (Li Shen and Mount Sinai (2013). GeneOverlap: Test and visualize gene overlaps. R package version 1.6.0.). Chip-Seq data has been submitted to NCBI’s Gene Expression Omnibus (GEO, <u>http://www.ncbi.nlm.nih.gov/geo/</u>); accession number GSE100440. Other statistical analyses were conducted using GraphPad Prism software, and values are presented as mean ± S.D. The statistical signficance of the differences between experimental groups was determined by two-tailed, unpaired Student’s t-test. Differences with a p-value of 0.05 were considered siginficant.

## References

1. Ehrt S & Schnappinger D (2009) Mycobacterial survival strategies in the phagosome: defence against host stresses. Cell Microbiol 11(8):1170–1178.

2. Sato K, Akaki T, & Tomioka H (1998) Differential potentiation of anti-mycobacterial activity and reactive nitrogen intermediate-producing ability of murine peritoneal macrophages activated by interferon-gamma (IFN-gamma) and tumour necrosis factor-alpha (TNF-alpha). Clin Exp Immunol 112(1):63–68.

3. Cooper AM, Segal BH, Frank AA, Holland SM, & Orme IM (2000) Transient loss of resistance to pulmonary tuberculosis in p47(phox-/-) mice. Infect Immun 68(3):1231–1234.

4. Lee PP, et al. (2008) Susceptibility to mycobacterial infections in children with X-linked chronic granulomatous disease: a review of 17 patients living in a region endemic for tuberculosis. Pediatr Infect Dis J 27(3):224–230.

5. Kumar A, et al. (2011) Redox homeostasis in mycobacteria: the key to tuberculosis control? Expert Rev Mol Med 13:e39.

6. Nambi S, et al. (2015) The Oxidative Stress Network of Mycobacterium tuberculosis Reveals Coordination between Radical Detoxification Systems. Cell Host Microbe 17(6):829–837.

7. Buchmeier NA, Newton GL, & Fahey RC (2006) A mycothiol synthase mutant of Mycobacterium tuberculosis has an altered thiol-disulfide content and limited tolerance to stress. J Bacteriol 188(17):6245–6252.

8. Saini V, et al. (2016) Ergothioneine Maintains Redox and Bioenergetic Homeostasis Essential for Drug Susceptibility and Virulence of Mycobacterium tuberculosis. Cell Rep 14(3):572–585.

9. Dillon SC & Dorman CJ (2010) Bacterial nucleoid-associated proteins, nucleoid structure and gene expression. Nat Rev Microbiol 8(3):185–195.

10. Martinez A & Kolter R (1997) Protection of DNA during oxidative stress by the nonspecific DNA-binding protein Dps. J Bacteriol 179(16):5188–5194.

11. Colangeli R, et al. (2009) The multifunctional histone-like protein Lsr2 protects mycobacteria against reactive oxygen intermediates. Proc Natl Acad Sci U S A 106(11):4414–4418.

12. Morikawa K, et al. (2006) Bacterial nucleoid dynamics: oxidative stress response in Staphylococcus aureus. Genes Cells 11(4):409–423.

13. Weinstein-Fischer D, Elgrably-Weiss M, & Altuvia S (2000) Escherichia coli response to hydrogen peroxide: a role for DNA supercoiling, topoisomerase I and Fis. Mol Microbiol 35(6):1413–1420.

14. Wang H, et al. (2012) Genetic and biochemical characteristics of the histone-like protein DR0199 in Deinococcus radiodurans. Microbiology 158(Pt 4):936–943.

15. Ko KC, Tai PC, & Derby CD (2012) Mechanisms of action of escapin, a bactericidal agent in the ink secretion of the sea hare Aplysia californica: rapid and long-lasting DNA condensation and involvement of the OxyR-regulated oxidative stress pathway. Antimicrob Agents Chemother 56(4):1725–1734.

16. Smith LJ, et al. (2010) Mycobacterium tuberculosis WhiB1 is an essential DNA-binding protein with a nitric oxide-sensitive iron-sulfur cluster. Biochem J 432(3):417–427.

17. Konar M, Alam MS, Arora C, & Agrawal P (2012) WhiB2/Rv3260c, a cell division-associated protein of Mycobacterium tuberculosis H37Rv, has properties of a chaperone. The FEBS journal 279(15):2781–2792.

18. Raghunand TR & Bishai WR (2006) Mycobacterium smegmatis whmD and its homologue Mycobacterium tuberculosis whiB2 are functionally equivalent. Microbiology 152(Pt 9):2735–2747.

19. Singh A, et al. (2007) Mycobacterium tuberculosis WhiB3 responds to O2 and nitric oxide via its [4Fe-4S] cluster and is essential for nutrient starvation survival. Proc Natl Acad Sci U S A 104(28):11562–11567.

20. Singh A, et al. (2009) Mycobacterium tuberculosis WhiB3 maintains redox homeostasis by regulating virulence lipid anabolism to modulate macrophage response. PLoS Pathog 5(8):e1000545.

21. Mehta M, Rajmani RS, & Singh A (2016) Mycobacterium tuberculosis WhiB3 Responds to Vacuolar pH-induced Changes in Mycothiol Redox Potential to Modulate Phagosomal Maturation and Virulence. J Biol Chem 291(6):2888–2903.

22. Chawla M, et al. (2012) Mycobacterium tuberculosis WhiB4 regulates oxidative stress response to modulate survival and dissemination in vivo. Mol Microbiol 85(6):1148–1165.

23. Mishra S, et al. (2017) Efficacy of beta-lactam/beta-lactamase inhibitor combination is linked to WhiB4-mediated changes in redox physiology of Mycobacterium tuberculosis. eLife 6.

24. Casonato S, et al. (2012) WhiB5, a transcriptional regulator that contributes to Mycobacterium tuberculosis virulence and reactivation. Infection and immunity 80(9):3132–3144.

25. Chen Z, et al. (2016) Mycobacterial WhiB6 Differentially Regulates ESX-1 and the Dos Regulon to Modulate Granuloma Formation and Virulence in Zebrafish. Cell reports 16(9):2512–2524.

26. Morris RP, et al. (2005) Ancestral antibiotic resistance in Mycobacterium tuberculosis. Proc Natl Acad Sci U S A 102(34):12200–12205.

27. Nandakumar M, Nathan C, & Rhee KY (2014) Isocitrate lyase mediates broad antibiotic tolerance in Mycobacterium tuberculosis. Nat Commun 5:4306.

28. Wolff KA, et al. (2015) A redox regulatory system critical for mycobacterial survival in macrophages and biofilm development. PLoS pathogens 11(4):e1004839.

29. Khan MZ, et al. (2017) Protein Kinase G confers survival advantage to Mycobacterium tuberculosis during latency like conditions. The Journal of biological chemistry.

30. Ohniwa RL, et al. (2006) Dynamic state of DNA topology is essential for genome condensation in bacteria. EMBO J 25(23):5591–5602.

31. Schneider R, Travers A, Kutateladze T, & Muskhelishvili G (1999) A DNA architectural protein couples cellular physiology and DNA topology in Escherichia coli. Mol Microbiol 34(5):953–964.

32. Master SS, et al. (2002) Oxidative stress response genes in Mycobacterium tuberculosis: role of ahpC in resistance to peroxynitrite and stage-specific survival in macrophages. Microbiology 148(Pt 10):3139–3144.

33. Hu Y & Coates AR (2009) Acute and persistent Mycobacterium tuberculosis infections depend on the thiol peroxidase TpX. PloS one 4(4):e5150.

34. Ung KS & Av-Gay Y (2006) Mycothiol-dependent mycobacterial response to oxidative stress. FEBS Lett 580(11):2712–2716.

35. Maksymiuk C, Balakrishnan A, Bryk R, Rhee KY, & Nathan CF (2015) E1 of alpha-ketoglutarate dehydrogenase defends Mycobacterium tuberculosis against glutamate anaplerosis and nitroxidative stress. Proc Natl Acad Sci U S A 112(43):E5834–5843.

36. Venugopal A, et al. (2011) Virulence of Mycobacterium tuberculosis depends on lipoamide dehydrogenase, a member of three multienzyme complexes. Cell Host Microbe 9(1):21–31.

37. Pathania R, Navani NK, Gardner AM, Gardner PR, & Dikshit KL (2002) Nitric oxide scavenging and detoxification by the Mycobacterium tuberculosis haemoglobin, HbN in Escherichia coli. Molecular microbiology 45(5):1303–1314.

38. Al-Attar S, et al. (2016) Cytochrome bd Displays Significant Quinol Peroxidase Activity. Scientific reports 6:27631.

39. Lu P, et al. (2015) The cytochrome bd-type quinol oxidase is important for survival of Mycobacterium smegmatis under peroxide and antibiotic-induced stress. Scientific reports 5:10333.

40. Fishbein S, van Wyk N, Warren RM, & Sampson SL (2015) Phylogeny to function: PE/PPE protein evolution and impact on Mycobacterium tuberculosis pathogenicity. Molecular microbiology 96(5):901–916.

41. Dame RT & Goosen N (2002) HU: promoting or counteracting DNA compaction? FEBS letters 529(2-3):151–156.

42. van Noort J, Verbrugge S, Goosen N, Dekker C, & Dame RT (2004) Dual architectural roles of HU: formation of flexible hinges and rigid filaments. Proceedings of the National Academy of Sciences of the United States of America 101(18):6969–6974.

43. Tessmer I, et al. (2005) AFM studies on the role of the protein RdgC in bacterial DNA recombination. Journal of molecular biology 350(2):254–262.

44. Kapuscinski J (1995) DAPI: a DNA-specific fluorescent probe. Biotechnic & histochemistry: official publication of the Biological Stain Commission 70(5):220–233.

45. Bhowmick T, et al. (2014) Targeting Mycobacterium tuberculosis nucleoid-associated protein HU with structure-based inhibitors. Nat Commun 5:4124.

46. Barksdale L & Kim KS (1977) Mycobacterium. Bacteriol Rev 41(1):217–372.

47. Takade A, Takeya K, Taniguchi H, & Mizuguchi Y (1983) Electron microscopic observations of cell division in Mycobacterium vaccae V1. J Gen Microbiol 129(7):2315–2320.

48. Vijay S, Nagaraja M, Sebastian J, & Ajitkumar P (2014) Asymmetric cell division in Mycobacterium tuberculosis and its unique features. Archives of microbiology 196(3):157–168.

49. Bhaskar A, et al. (2014) Reengineering redox sensitive GFP to measure mycothiol redox potential of Mycobacterium tuberculosis during infection. PLoS Pathog 10(1):e1003902.

50. Vilcheze C, et al. (2008) Mycothiol biosynthesis is essential for ethionamide susceptibility in Mycobacterium tuberculosis. Mol Microbiol 69(5):1316–1329.

51. Buchmeier NA, Newton GL, Koledin T, & Fahey RC (2003) Association of mycothiol with protection of Mycobacterium tuberculosis from toxic oxidants and antibiotics. Mol Microbiol 47(6):1723–1732.

52. Wu ML, et al. (2016) Developmental transcriptome of resting cell formation in Mycobacterium smegmatis. BMC genomics 17(1):837.

53. Rustad TR, Harrell MI, Liao R, & Sherman DR (2008) The enduring hypoxic response of Mycobacterium tuberculosis. PLoS One 3(1):e1502.

54. Galagan JE, et al. (2013) The Mycobacterium tuberculosis regulatory network and hypoxia. Nature 499(7457):178–183.

55. Kahramanoglou C, et al. (2014) Genomic mapping of cAMP receptor protein (CRP Mt) in Mycobacterium tuberculosis: relation to transcriptional start sites and the role of CRPMt as a transcription factor. Nucleic acids research 42(13):8320–8329.

56. Field Y, et al. (2008) Distinct modes of regulation by chromatin encoded through nucleosome positioning signals. PLoS Comput Biol 4(11):e1000216.

57. Prieto AI, et al. (2012) Genomic analysis of DNA binding and gene regulation by homologous nucleoid-associated proteins IHF and HU in Escherichia coli K12. Nucleic Acids Res 40(8):3524–3537.

58. Cole ST, et al. (1998) Deciphering the biology of Mycobacterium tuberculosis from the complete genome sequence. Nature 393(6685):537–544.

59. Lew JM, Kapopoulou A, Jones LM, & Cole ST (2011) TubercuList--10 years after. Tuberculosis 91(1):1–7.

60. Wu J, et al. (2017) WhiB4 Regulates the PE/PPE Gene Family and is Essential for Virulence of Mycobacterium marinum. Scientific reports 7(1):3007.

61. Cho BK, Knight EM, Barrett CL, & Palsson BO (2008) Genome-wide analysis of Fis binding in Escherichia coli indicates a causative role for A-/AT-tracts. Genome research 18(6):900–910.

62. Grainger DC, Hurd D, Harrison M, Holdstock J, & Busby SJ (2005) Studies of the distribution of Escherichia coli cAMP-receptor protein and RNA polymerase along the E. coli chromosome. Proceedings of the National Academy of Sciences of the United States of America 102(49):17693–17698.

63. Kahramanoglou C, et al. (2011) Direct and indirect effects of H-NS and Fis on global gene expression control in Escherichia coli. Nucleic Acids Res 39(6):2073–2091.

64. Burian J, Ramon-Garcia S, Howes CG, & Thompson CJ (2012) WhiB7, a transcriptional activator that coordinates physiology with intrinsic drug resistance in Mycobacterium tuberculosis. Expert Rev Anti Infect Ther 10(9):1037–1047.

65. Bush MJ, Chandra G, Bibb MJ, Findlay KC, & Buttner MJ (2016) Genome-Wide Chromatin Immunoprecipitation Sequencing Analysis Shows that WhiB Is a Transcription Factor That Cocontrols Its Regulon with WhiA To Initiate Developmental Cell Division in Streptomyces. MBio 7(2):e00523–00516.

66. Facey PD, et al. (2009) Streptomyces coelicolor Dps-like proteins: differential dual roles in response to stress during vegetative growth and in nucleoid condensation during reproductive cell division. Mol Microbiol 73(6):1186–1202.

67. Ouafa ZA, Reverchon S, Lautier T, Muskhelishvili G, & Nasser W (2012) The nucleoid-associated proteins H-NS and FIS modulate the DNA supercoiling response of the pel genes, the major virulence factors in the plant pathogen bacterium Dickeya dadantii. Nucleic Acids Res 40(10):4306–4319.

68. Ushijima Y, et al. (2014) Nucleoid compaction by MrgA(Asp56Ala/Glu60Ala) does not contribute to staphylococcal cell survival against oxidative stress and phagocytic killing by macrophages. FEMS Microbiol Lett 360(2):144–151.

69. Bartek IL, et al. (2014) Mycobacterium tuberculosis Lsr2 is a global transcriptional regulator required for adaptation to changing oxygen levels and virulence. MBio 5(3):e01106–01114.

70. Dwyer DJ, Camacho DM, Kohanski MA, Callura JM, & Collins JJ (2012) Antibiotic-induced bacterial cell death exhibits physiological and biochemical hallmarks of apoptosis. Mol Cell 46(5):561–572.

71. Wu ML, Gengenbacher M, & Dick T (2016) Mild Nutrient Starvation Triggers the Development of a Small-Cell Survival Morphotype in Mycobacteria. Front Microbiol 7:947.

72. Vijay S, Anand D, & Ajitkumar P (2012) Unveiling unusual features of formation of septal partition and constriction in mycobacteria--an ultrastructural study. Journal of bacteriology 194(3):702–707.

## References

1. Chawla M, et al. (2012) Mycobacterium tuberculosis WhiB4 regulates oxidative stress response to modulate survival and dissemination in vivo. Mol Microbiol 85(6):1148–1165.

2. Venkataraman B, Vasudevan M, & Gupta A (2014) A new microarray platform for whole-genome expression profiling of Mycobacterium tuberculosis. J Microbiol Methods 97:34–43.

3. Kahramanoglou C, et al. (2011) Direct and indirect effects of H-NS and Fis on global gene expression control in Escherichia coli. Nucleic Acids Res 39(6):2073–2091.

4. Kahramanoglou C, et al. (2014) Genomic mapping of cAMP receptor protein (CRP Mt) in Mycobacterium tuberculosis: relation to transcriptional start sites and the role of CRPMt as a transcription factor. Nucleic acids research 42(13):8320–8329.

